# Archaeal SegAB forms a bipolar structure that promotes chromosome segregation in spherical cells

**DOI:** 10.1101/2025.04.15.649018

**Authors:** Arthur Charles-Orszag, Samuel J Lord, Nadia Herrera, Luke Strauskulage, Arghya Bhowmick, Tom Goddard, Bianca Wassmer, Marleen van Wolferen, Garrison Asper, Amy Flis, Johnny Rodriguez, Sy Redding, Oren Rosenberg, Sonja-Verena Albers, R. Dyche Mullins

**Author notes:** Correspondence to R. Dyche Mullins.

## Abstract

Archaeal *segAB* operons are thought to promote chromosome segregation, but their mechanism remains unknown. We employ comparative genomics, structural biology, genetic knockouts, and quantitative cell biology to investigate how SegA and SegB proteins work together to segregate chromosomes in the thermophilic archaeon *Sulfolobus acidocaldarius*. *In vitro*, SegB binds a centromeric DNA sequence adjacent to the *segAB* operon, and *in vivo* forms a distinct focus on each segregating chromosome. SegA, a ParA-like ATPase, binds DNA non-specifically *in vitro* and promotes chromosome compaction and segregation *in vivo*. During division, SegA shifts from chromosome-associated puncta to form a single, elongated figure that runs between separating SegB foci. Late in division, SegA retreats to regions surrounding separated SegB foci. Elongated SegA figures appear in *segB* knockout cells but no longer lie perpendicular to the division plane. We propose that SegA and SegB interact to form a bipolar, DNA-segregating structure radically different from bacterial ParABS systems.

## Introduction

How do dividing cells distribute genetic material to their daughters? In eukaryotes, the microtubule-based mitotic spindle captures, aligns, and segregates chromosomes (Valdez et al., 2023). In bacteria and archaea, the situation is less clear. Many bacterial (and some archaeal) chromosomes contain partitioning (Par) operons that promote proper chromosome segregation. The most common partitioning operons in both bacteria and archaea rely on ParA-type ATPases (Merino-Salomón, Babl, and Schwille 2021; Pulianmackal et al. 2023). Work on ParA-based systems has yielded attractive models for bacterial DNA segregation, but archaeal variants remain mysterious (Lim et al. 2014).

ParA homologs are best understood in bacteria, where many of them function as components of ParABS segregation systems. These bacterial ParABS systems comprise (i) a centromeric DNA locus, *parS*; (ii) a centromere-binding protein, ParB; and (iii) an ATPase, ParA (Mohl and Gober 1997; Toro et al. 2008). The ParB protein forms a dimeric clamp that loads onto DNA specifically at the *parS* locus (Jalal et al. 2021; Osorio-Valeriano et al. 2019). The ParA protein binds nonspecifically to DNA, with an affinity regulated by its ATPase activity. In most species studied to date, ATP-ParA binds DNA more tightly than the ADP-bound form (Lim et al. 2014), and ParB promotes ParA dissociation by accelerating ATP hydrolysis (Easter and Gober 2002; Lim et al. 2014).

Recent models based on data from rod-shaped bacteria argue that ParABS promotes DNA segregation via a reaction-diffusion process. In these models, ParB/*parS* segregation complexes create a gradient of DNA-bound ParA by locally promoting ATP hydrolysis and dissociation from DNA (Chin Hwang et al. 2012). Experimentally, ParA disappears from regions surrounding ParB/*parS* complexes even before initiation of chromosome segregation is detectable (Surovtsev, Lim, and Jacobs-Wagner 2016). Once formed, the self-generated ParA gradient is thought to rectify random thermal motion of a mobile ParB/*parS* segregation complex, driving it on a journey from one cell pole to the other. In a version of this model called the DNA relay, ParB/*parS* movement is further assisted by the spring-like mechanics of chromosomal DNA (Lim et al. 2014). During this trip, the ParA gradient retreats toward the distal cell pole, just ahead of the steadily advancing ParB/*parS* complex. This picture is way less clear in non-rod-shaped bacteria, and only recently was it reported that *Neisseria gonorrhoeae* ParB does not localize to the cell poles but rather aligns with the long axis of the cell during cytokinesis (Bandekar et al., 2025).

Some archaeal chromosomes contain genes encoding ParA-like proteins, but little is known about their function. Genomes of both *Saccharolobus solfataricus* (*Sso*) and *Sulfolobus acidocaldarius* (*Saci*), for example, encode an ATPase called SegA that shares ∼34% sequence identity with bacterial ParA proteins. The gene immediately adjacent to *segA*, however, exhibits no sequence similarity to bacterial *parB*, and encodes an archaea-specific protein called SegB (Kalliomaa-Sanford et al. 2012). Both SegA and SegB are expressed in a cell cycle-dependent manner (Lundgren and Bernander, 2007), and overexpression or deletion of the *segAB* operon in both *Sso* and *Saci* produce growth defects, abnormal nucleoid morphology, and anucleate cells, consistent with a role in chromosome segregation (Kalliomaa-Sanford et al. 2012; Kabli et al. 2025). Additionally, *Sso*—but not *Saci*—encodes a second NTPase, SegC, that co-assembles into filaments with SegA and DNA *in vitro* (Lin et al., 2024). Structurally, the SegA protein from *Sso* resembles bacterial ParA, but with a variant dimer architecture (Yen et al. 2021), while the enigmatic SegC adopts a Thiamin diphosphate-binding fold (Lin et al. 2024). Unfortunately, only the C-terminal two thirds of the *Sso* SegB protein (residues 34-109) has been crystallized. This truncated SegB adopts a helix-turn-helix fold, forms dimers, and binds DNA—but it lacks the SegA-interacting N-terminal region (Yen et al. 2021).

Sequence analysis and comparative genomics reveal a correlation between the presence of a *segB* gene and spherical cell shape, suggesting that SegB evolved to help break cellular symmetry and establish an axis for chromosome segregation in non-rod-shaped archaea. But until now, no clear models have emerged for how SegA and SegB promote chromosome segregation. SegB binds multiple loci throughout the *Sso* chromosome, including the centromeric DNA sequence, *segS*, located upstream of the *segAB* operon (Kabli et al. 2025; Kalliomaa-Sanford et al. 2012; Yen et al. 2021). SegB in complex with DNA was proposed to form a superhelix that links distant chromosomal loci to form discrete SegB/DNA clusters throughout the cytoplasm of non-dividing cells, and to promote ATP hydrolysis by SegA (Kalliomaa-Sanford et al., 2012). Similar to bacterial ParA, SegA binds non-specifically to DNA but, unlike in bacteria, this binding does not appear to be regulated by ATP hydrolysis (Yen et al. 2021). First thought to assemble into short polymers (Kalliomaa-Sanford et al. 2012), SegA was later reported to form discrete foci in non-dividing cells, leading to the hypothesis that SegA and SegB combine genome organization and chromosome segregation activities (Kabli et al., 2025).

We chose to study the SegAB system in *Saci*, in part because the lack of SegC suggested that its mechanism might be simpler than that of *Sso*. We first crystallized full-length *Saci* SegB and solved its structure, which revealed a similar helix-turn-helix motif but a very different dimer architecture from that reported by Yen et al. (2021) for the truncated *Sso* SegB. *In vitro*, we find that SegB from *Saci* binds a specific sequence (*segS*), immediately upstream of the *segAB* operon and close to one of the three origins of replication. In non-dividing cells, SegB is minimally expressed and exhibits no clear localization pattern, but during division it forms two well defined chromosome-associated foci that eventually localize to opposite poles of the cell. In *segA*-knockout cells, SegB forms several foci, suggesting that SegA acts to limit or concentrate its localization.

SegA undergoes a more dramatic rearrangement during division. Mostly absent in non-dividing cells, SegA appears as chromosome-associated foci in early stages of division, but then forms an elongated, filament-like figure running between SegB foci as division progresses. In wildtype cells, this divisional SegA figure always runs perpendicular to the plane of cell division. Loss of SegB does not abrogate formation of this SegA figure, but randomizes its orientation with respect to the division plane. From these data, we propose a model for SegAB-mediated chromosome segregation completely distinct from the current understanding of the ParABS mechanism in bacteria.

## Results

### The *S. acidocaldarius* chromosome contains a *segAB* operon that promotes chromosome segregation

Genome analysis reveals that *S. acidocaldarius* (*Saci*) encodes homologs of both *segA* and *segB* genes (saci0204 and saci0203), originally described in *Sa. solfataricus* (*Sso*). The SegA and SegB proteins encoded by these *Saci* genes share 71% and 53% sequence identity with their *Sso* counterparts. Similar to Sso, the 3’ region of the *Saci segA* gene overlaps the 5’ region of *segB* (Kalliomaa-Sanford et al., 2012), suggesting that the two genes are transcribed as a single unit and likely function together in the same process (Wurtzel et al., 2010). The genomic context of the *segAB* region of Saci differs significantly from that of *Sso* (Fig. 1a and 3c), indicating that *segA* and *segB* have maintained close proximity across evolutionary time scales and/or have moved together as a unit during lateral gene transfer events. While the SegA proteins from *Saci* and *Sso* are of similar size, the Saci SegB protein is almost 50% larger than the *Sso* version (151 versus 109 amino acids) (Fig. 1a and 3c). This difference illustrates the size range of archaeal SegB proteins, which can vary more than five-fold in length (Kabli et al., 2025; Kalliomaa-Sanford et al., 2012) from species to species (see below).

**Figure 1.**
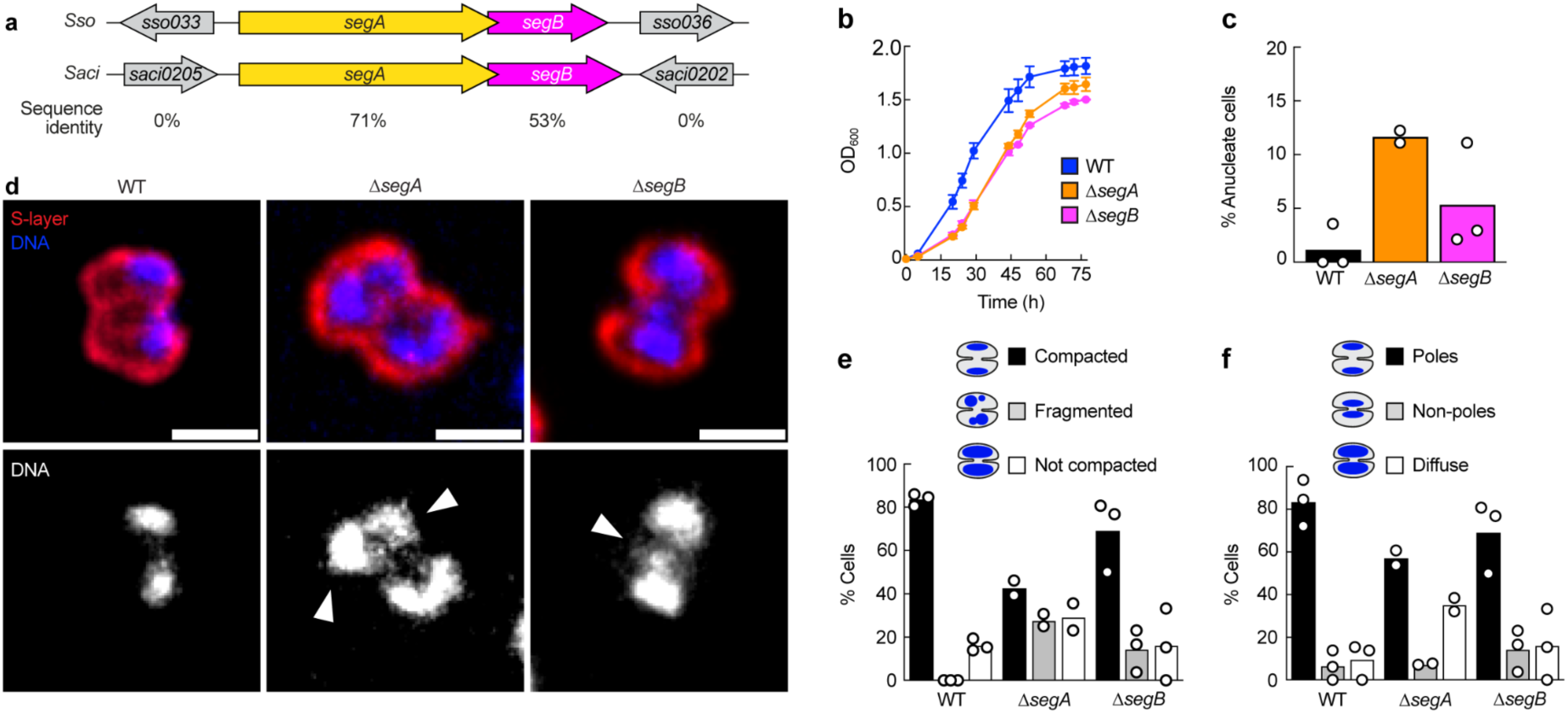
*S. acidocaldarius* SegAB promotes chromosome segregation and compaction. **a.** Comparison between the *segAB* operons of *Saccharolobus solfataricus* (Sso) and *S. acidocaldarius* (Saci). **b.** Growth of Δ*segA* and Δ*segB* deletion strains compared to WT. Data points represent the mean ±SEM of three technical replicates; representative of three biological replicates. **c.** Quantification of the absence of chromosomal DNA (anucleate) in WT, Δ*segA* and Δ*segB* deletion strains. **d.** Super-resolution fluorescence of chromosomal DNA in indicated strains. Arrowheads indicate fragmented chromosomes. Scale bars, 1 µm. **e, f.** Quantitative analysis of the aspect and localization of chromosomal DNA in WT, Δ*segA* and Δ*segB* cells. A total of 161 (WT), 81 (Δ*segA*) and 109 (Δ*segB*) cells were analyzed; each dot represents the mean of a biological replicate.

To understand the function of the *segAB* operon in *Saci* we created deletion strains for both *segA* and *segB*. Although viable, both mutant strains exhibited a significant growth defect compared to WT cells (Fig. 1b). Consistent with results from *Sso* (Kabli et al., 2025; Kalliomaa-Sanford et al., 2012), we also observed an increase in the number of anucleate cells in both knockout strains: increasing from 1.19%±2.06, in wildtype cells, to 11.68%±0.80 and 5.39%±4.9, in *segA* and *segB* knockouts respectively (Fig. 1c). To compare chromosome morphology and localization in wildtype and mutant strains, we fixed cells after covalently labeling the S-layer, stained DNA with DAPI, and imaged them by super-resolution (iSIM) light microscopy (Fig. 1d). We scored only cells undergoing division, as judged by asymmetric cell shape and the presence of detectable ingression of a cleavage furrow. To avoid observer bias we performed imaging and quantification double-blind. Chromosomes more often appeared fragmented in both *segA* and *segB* deletion strains (Fig. 1e, left graph). Interestingly, knocking out *segA* caused a bigger increase in the fraction of cells with uncompacted and diffusely localized chromosomes (Fig. 1e) than loss of *segB*. From these experiments we conclude that SegA and SegB promote chromosome segregation in *Saci*, and that SegA plays an additional role in compaction of segregating chromosomes.

Finally, we used high-temperature (76°C) time-lapse imaging to compare cytokinesis in wildtype and mutant cells. When we tracked visible cleavage furrow ingression, we saw no difference in the rate or duration of cytokinesis in the Δ*segB* strain compared with wildtype (Fig. S1 c, d). When we used mother-daughter cell separation as a marker for abscission, however, we noted a slight (statistically non-significant) delay in Δ*segB* cells (Fig. S1 e, f).

Sequence-specific binding of SegB to sites in the *S. acidocaldarius* chromosome

For *in vitro* experiments, we expressed the full-length SegB protein fused to a SUMO (Small Ubiquitin-like Modifier) tag. Following purification, we cleaved and separated the SUMO tag to produce SegB with no additional amino acid residues on either the N- or C-terminus. The mobility of this purified SegB in gel electrophoresis (SDS-PAGE) assays was consistent with a molecular weight of 20 kDa, slightly higher than its predicted molecular weight of 17.4 kDa (Fig. S2). In addition to this wildtype SegB, we also created a fluorescent SegB derivative by mutating a surface-exposed serine residue (S56) to cysteine (SegB_S56C_) and then reacting this mutant protein with a maleimide-functionalized fluorescent dye (AlexaFluor 568).

We used purified SegB and electrophoretic mobility shift assays (EMSAs) to look for specific binding sequences in the chromosomal region near the *segAB* operon (Fig. 2a and Fig. S2). SegB binds strongly to a 300-bp fragment immediately upstream of *segAB* but only weakly interacts with DNA oligonucleotides from the 3’ flanking region (Fig. S2). We narrowed down the SegB binding site by splitting the proximal 100 base pairs on the 5’ side of the *segA* start codon into 20-bp DNA fragments with 5-nucleotide overlaps (Fig. 2a). In these experiments SegB bound specifically to the two DNA fragments closest to *segA* start codon. From this we deduce that SegB binds specifically to the DNA sequence that runs from nucleotides -25 to -6 on the 5’ side of the *segA* start codon (5’-AGGGTATAGAGTTAAGAGT-3’). Consistent with established nomenclature (Kalliomaa-Sanford et al., 2012), we refer to this sequence as the *Saci segS* motif. SegB-*segS* binding was sequence-specific as judged by EMSA assays performed using SegB and 20-bp DNA molecules containing either a native or scrambled *segS* sequence (Fig. 2c).

**Figure 2.**
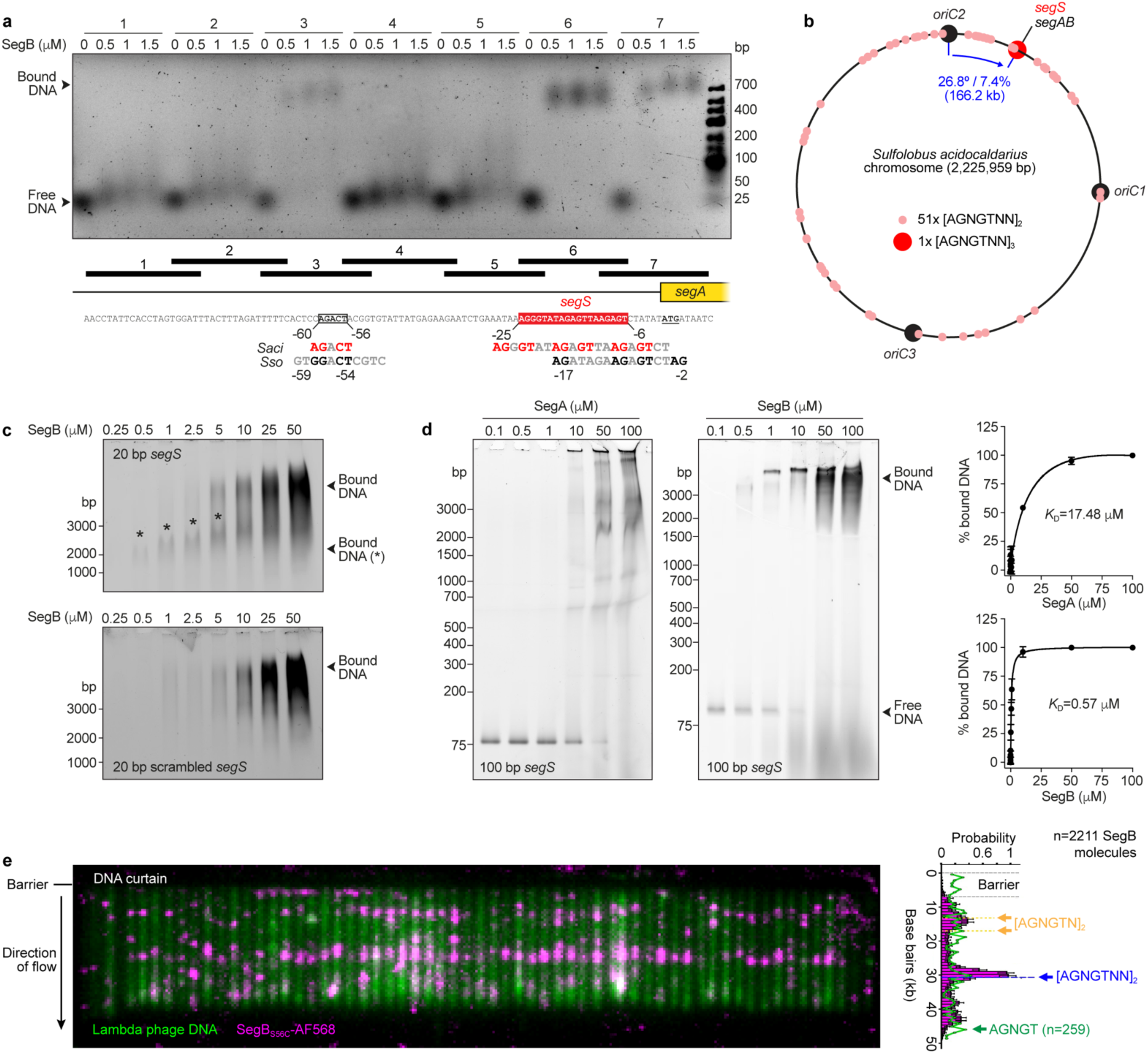
*Sulfolobus acidocaldarius* SegB binds centromeric DNA sequence, *segS*. **a.** Identification of the centromeric sequence and SegB binding partner *segS* upstream of the *segAB* operon by EMSA. Alignment with the corresponding region in Sso shows partial conservation of *segS*. **b.** Bioinformatic mapping of one full *segS* sequence (containing three repeats) and 51 shorter *segS*-like motifs onto Saci chromosome. **c.** EMSA showing binding of SegB to *segS* and scrambled DNA sequences. Asterisks indicate *segS*-specific binding. **d.** EMSA showing binding of SegA and SegB to a 100-bp DNA fragment corresponding to the 50-bp upstream and downstream of the *segAB* operon containing *segS*. The graphs represent the determination of SegA-DNA and SegB-DNA respective dissociation constants by densitometry in two independent experiments each. **e.** DNA curtain assay showing the binding of single molecules of SegB_S56C_ labeled with AlexaFluor 568 to lambda phage DNA under flow. The bar plot shows the distribution of all SegB_S56C_ molecules imaged along lambda phage DNA. Single, double and near-double *segS* motifs present in the DNA are indicated. The highest binding peak corresponds to a double repeat of the *segS* motif.

Unlike SegB, recombinant SegA binds DNA with no obvious sequence specificity (Fig. 2d and Fig. S2). By EMSA SegA binds with lower affinity (*K*_d_=17.48 µM) than SegB (*K*_d_=0.57 µM) to a 100-bp DNA fragment containing the 50 base pairs upstream and downstream of *segAB* (Fig. 2d).

*Saci segS* exhibits weak homology to the *segS* motif found in Sso (Fig. 2a) but, unlike the Sso version, it does not form a pseudo-palindrome. Instead, *Saci segS* comprises three repeats of the motif [AGNGTNN]. Interestingly, SegB also bound (although with lower apparent affinity) to a third DNA fragment located -60 to -56 base pairs upstream of *segA* start codon. This oligonucleotide contains an AGACT sequence reminiscent of the three [AGNGTNN] repeats found in *segS*. The full-length *segS* (i.e. the triple repeat [AGNGTNN]_3_) occurs only once in the entire *Saci* chromosome, adjacent to the *segAB* operon and 166.2 kb (7.6% of the genome size) from *oriC2*, one of three origins of replications in *Saci* (Fig. 2b). For comparison, ∼92% of bacterial *parS* sites fall within 15% of the chromosome length from *oriC*, the origin of replication in primary bacterial chromosomes (Livny et al., 2007). Because we found that SegB can bind partial *segS* motifs *in vitro* (Fig. 2c), we looked for additional incomplete *segS* motifs in the *Saci* genome and found 51 double repeats ([AGNGTNN]_2_) scattered across the chromosome (Fig. 2b).

To assess the ability of SegB to bind partial and/or imperfect *segS* sequences, we employed a single-molecule DNA curtain approach (Fig. 2e; Greene et al., 2010). Briefly, we immobilized ∼50 kbp molecules of double-stranded DNA from bacteriophage λ in a microfluidic flow chamber, via a supported lipid bilayer. We labeled the DNA molecules with the intercalating dye YOYO-1. We then pulsed fluorescently labeled SegB protein into the flow chamber and imaged both DNA and SegB by Total Internal Reflection Fluorescence (TIRF) microscopy. In this assay labeled SegB binds predominantly to two regions in the lambda phage DNA, located at ∼15 kb and ∼30 kb positions, with probabilities of ∼40% and ∼80% respectively (Fig. 2e). Little or no binding was observed outside of these loci, despite the presence of 259 AGNGT motifs separated by more than one base pair. Interestingly, the region around 15 kb contains two [AGNGTN]_2_ motifs, corresponding to two *segS*-like sequences where two AGNGT sequences are separated by one base pair instead of two, while the region around 30 kb contains one true [AGNGTNN]_2_ *segS*-like sequence as found throughout Saci chromosome (Fig. 2b).

Together, our DNA-protein interaction studies demonstrate that *Saci* SegA binds DNA non-specifically, with potential to interact throughout the entire chromosome. In contrast, SegB exhibits both sequence-specific and non-specific DNA binding. The affinity of SegB for segS-like sequences composed of partial and/or imperfect repeat motifs suggests that SegB may bind multiple partial and/or imperfect *segS* sites across the chromosome in addition to the high-affinity *segS* site proximal to the *segAB* operon.

### Structure of the *S. acidocaldarius* SegB dimer

SegB from *Saci* is considerably larger than its counterpart in *Sso* (151 versus 109 amino acids), with most of the difference down to two sequence insertions. One 30-residue insertion appears at a position equivalent to R12/K13 in *Sso* SegB and another, 12-residue insertion, occurs at N26/A27 of *Sso* SegB. Bioinformatic analyses fail to predict secondary structure for these insertions and their potential function is unknown. In addition, the previously determined structure of *Sso* SegB (Yen et al. 2021) is missing the N-terminal one third of the molecule (residues 1-33), which contains the SegA-interacting region. To fill these gaps, we crystallized full-length *Saci* SegB and solved its atomic structure at 1.85Å resolution (Fig. 3a). Consistent with previous structural work (Kalliomaa-Sanford et al., 2012; Yen et al., 2021) and our circular dichroism measurements (Fig. S1), SegB forms a dimer (Fig. 3a) of globular proteins, composed mostly of ɑ-helices with a small percentage of β-strand and disordered regions (Fig. 3a). Comparing *Sso* and *Saci* structures reveals that all secondary structural elements of Sso SegB also appear in the *Saci* protein, namely the five alpha helices (ɑ1 to ɑ5) and the lone beta strand (β2).

**Figure 3.**
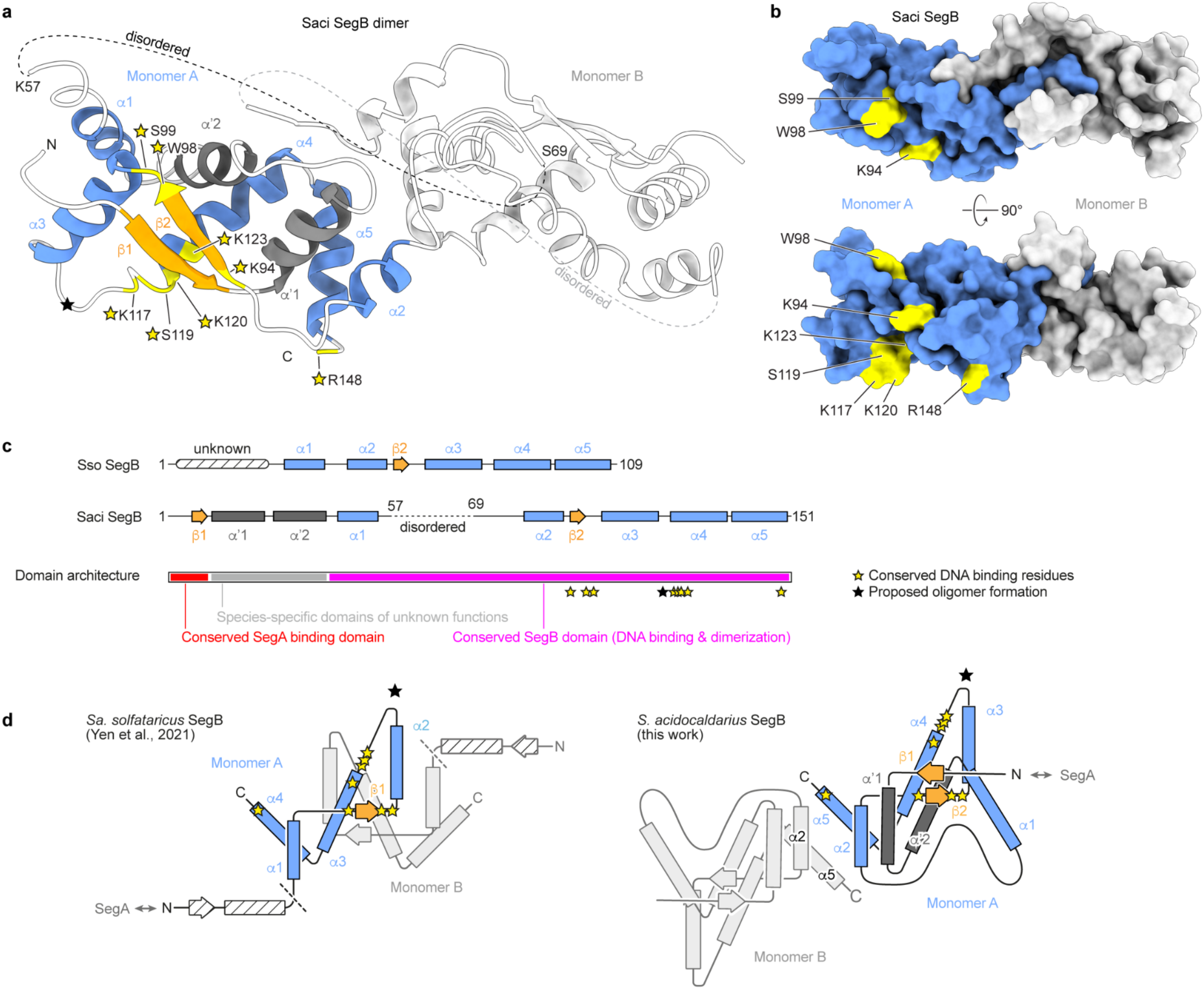
Structure determination of SegB in *Sulfolobus acidocaldarius*. **a.** Ribbon diagram of Saci SegB dimer. In monomer A, conserved alpha helices are colored in blue while helices specific to Saci are colored in dark gray. Conserved beta strands are colored in gold. Gold stars indicate conserved residues involved in DNA binding, while the light grey star corresponds to the residue proposed to be involved in oligomer formation in Sso SegB. Monomer B is colored in light grey for clarity. **b.** Surface representation of Saci SegB dimer showing the head-to-tail dimer conformation and solvent accessibility of all the conserved residues involved in DNA binding. **c.** Secondary structure and domain organization of Saci and Sso SegB homologs. **d.** Comparison between the folding of truncated Sso SegB and full-length Saci SegB.

The conserved N-terminal SegA-binding domain (Yen et al., 2021)—missing from the *Sso* SegB structure—contains an additional beta strand (β1), which contacts β2 to form an antiparallel sheet (Fig. 3a, d). While the region between residues K57 and S69 did not form a stable tertiary structure, the 30-residue insertion unique to *Saci* SegB forms two alpha helices (ɑ’1 and ɑ’2), arranged in a helix-turn-helix configuration. This arrangement enables β1 and β2 to interact while preserving the conserved ribbon-helix-helix motif formed by β2, ɑ3 and ɑ4 (β1, ɑ2, ɑ3 in *Sso* SegB, (Yen et al., 2021)) (Fig. 3c, d). All eight conserved residues involved in DNA binding, K94, W98, S99, K117, S119, K120, K123 and R148 (corresponding to *Sso* K52, W56, S57, K75, S77, K78, K81, R106), as well as residues proposed to be involved in oligomerization (Yen et al., 2021) are exposed on the surface of the protein (Fig. 3a, b).

Surprisingly, the dimer architecture of *Saci* SegB differs strikingly from that reported for the *Sso* protein (Fig. 3a, b, d). In SegB from *Sso*, dimerization involves the β1 strand from each monomer, interacting to form an antiparallel sheet (Fig. 3d) (Yen et al., 2021). In our *Saci* SegB structure, two beta strands from the same polypeptide (β1 and β2) form an *intra*-molecular antiparallel sheet (Fig.3 a, d) similar to the *inter*-molecular interaction reported for *Sso* SegB. The dimer interface in our structure is formed by the ɑ2 and ɑ5 helices from each monomer (Fig. 3 a, d). To verify that our structure for the *intra*-molecular beta sheet is correct, we inspected the electron density of side chains in this region. Electron density maps showed clearly that the sequences of the two interacting beta strands are different, representing β1 and β2 from the same polypeptide, rather two β1 strands from different polypeptides (Fig. S3). Since the residues forming β1 in our structure were missing from the truncated *Sso* SegB (Yen et al., 2021) we suggest that the N-terminal region of *Sso* SegB may also contain an additional beta strand (Fig. 3d), and that full-length SegB from *Sso* may also form a dimer similar to the one in *Saci*. In any case, it is remarkable that in *Saci* SegB the central helix-turn-helix fold and the solvent accessibility of the residues involved in binding to DNA are conserved despite the evolution of two additional non-conserved N-terminal alpha helices.

### Sequence diversity and evolutionary history of archaeal SegB proteins

The size difference between *Saci* and *Sso* SegB proteins inspired us to compare more SegB homologs. We used the *Sso* SegB protein sequence as a query to the Basic Local Alignment Search Tool (BLAST; (Altschul et al., 1990)), searching against the entire NCBI non-redundant sequence database (see Materials and Methods). The initial search returned 117 hits, but we winnowed this list by removing redundant or incomplete sequences as well as sequences from unidentified or poorly annotated species. We kept only sequences containing the signature, C-terminal SegB motif (K/R-D/N-P-K/R-I-G/T-V/L-WS-Y/P-P/E), which left 72 unique sequences from well-defined species (Fig. S4). All these sequences were adjacent to a predicted ParA-like ATPase. Predicted SegB homologs fell into two clusters: a large group (50 sequences) in the Thermoproteota and a smaller group (22 sequences) in the Euryarchaeota. Thermoproteota sequences clustered into eight further subgroups, with pairwise identities to *Sso* SegB as low as 39% (in *Sulfodiicoccus acidiphilus*). Euryarchaeota sequences fell into four subgroups (Fig. S4a) with generally lower pairwise identities to *Sso* SegB (from 30% in *Methanosphaera sp.* to 49% in *Thermococcus aggregans*). The predicted length of SegB proteins varied from 68 to 331 amino acids, with most homologs falling between 105-120 amino acids (Fig. S4b). The low degree of conservation among SegB homologs largely reflects variability in the middle of the polypeptides, a region that contains most of the species-specific features (Fig. S4c). In contrast, the first 10-12 and last 60 amino acids are highly conserved across the entire SegB family. These results suggest an unusual evolutionary trajectory of SegB homologs.

Within the variable nucleotide sequences in the middle regions of the *segB* genes we noticed an enrichment of G[A]_n_ tracts of various lengths. Such sequences—for example AAGAAGAAAAAGAAAAAG (Fig. 4a)—can form genomically unstable regions, prone to slipped-strand mispairing. Because SegB homologs with sequences longer than ∼105-120 amino acids all contain insertions at this site, we propose that the long SegB homologs arose from a short ancestor via progressive sequence expansion at the G[A]_n_ tracts. The SegB homologs of Metallosphaera species represent an extreme example of this sequence expansion. These homologs have 82-182 additional amino acids forming domains of unknown function. These domains consist of 5-13 repeats of a Metallosphaera-specific 14-amino acid motif located between the conserved SegA-binding and DNA-binding domains (Fig. 4a). The longer homologs likely evolved from a shorter ancestor similar to the short homologs found in other species of this family. Interestingly, in *M. javensis* and *M. sedula*, our search revealed that multiple short and long copies of SegB may co-exist in the same species (Fig. S5).

**Figure 4.**
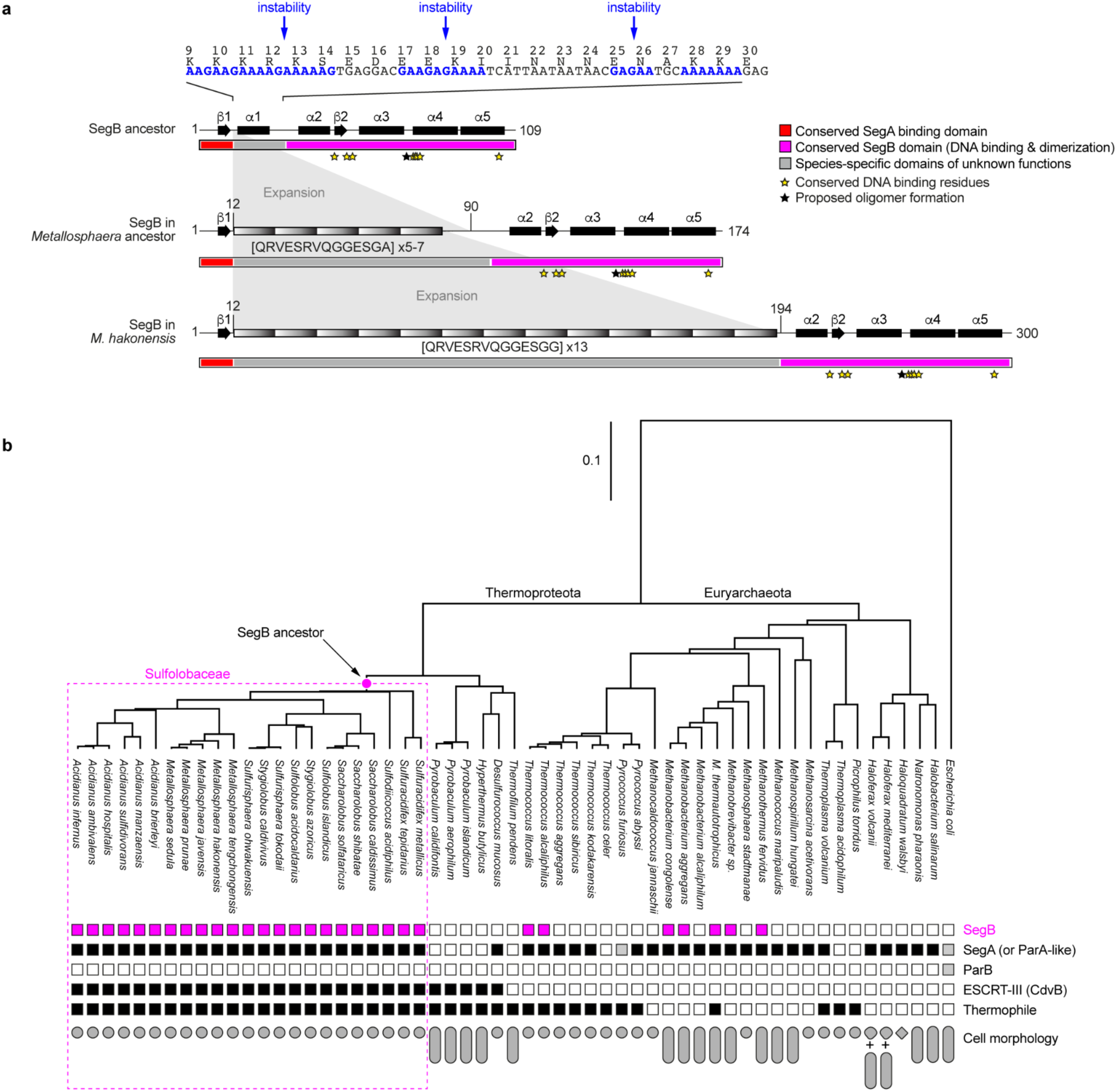
Evolution of SegB and shared characteristics of SegB-encoding archaea. **a.** Sites of species-specific domain insertions correspond to loci in *segB* 5’ that contain GA and GAA repeats. The proposed SegB ancestor corresponds to a *Sso*-like short SegB. Insertion of additional domains at these sites in a species-specific manner may have led to longer SegB homologs in extant species such as in *M. hakonensis*. Conserved domains correspond to the C- and N-terminal parts of SegB and contain DNA- and SegA-binding residues. **b.** Comparative genomics suggests that SegB is involved in cell polarity in non-rod-shaped Thermoproteota. Compared to other groups, all SegB-encoding Thermoproteaota have in common to be non-rod-shaped thermophiles that seemingly rely on ESCRT-III for cell division and don’t possess a homolog of bacterial ParB, suggesting that SegB is involved in the establishment or the interpretation of polarity cues in non-rod-shaped cells, in lieu of ParB which assumes a similar function in rod-shaped bacteria.

Mapping the length of all SegBs onto the natural phylogenetic tree of representative Thermoproteota and Euryarchaeota species strongly suggested that the same expansion mechanism can explain the lengths of all predicted SegB homologs (Fig. 4a). Since the majority of SegB sequences contain few (or no) extra domains between the conserved N- and C-terminal domains, the common ancestor of all known SegBs was short. SegB is present in all Sulfolobaceae, but absent from the rest of the Thermoproteota (e.g. *Pyrobaculum* or *Desulfurococcus*), suggesting that SegB first appeared in the common ancestor of Sulfolobaceae (Fig. 4b, Fig. S5). In eight families of Sulfolobaceae, however, SegB homologs exhibit lineage-specific insertions of additional domains of comparable lengths between the conserved N- and C-terminal domains, likely corresponding to eight independent expansion events that yielded SegB homologs with a high level of conservation among closely related species but a relatively lower level of conservation overall (Fig. S5). The scarcity and low level of conservation among SegB homologs in Euryarchaeota (especially in the conserved N-terminal domain) suggests that these homologs were likely acquired by horizontal gene transfer from Sulfolobaceae. More specifically, we propose that SegB was transferred twice: once in the ancestor of *Thermococcus alcaliphilus* and *T. litoralis*, and once in the ancestor of Methanobacteriales. These homologs evolved following a similar expansion mechanism, yielding longer homologs containing lineage-specific N-terminal domains of unknown function, or alternatively were lost (Fig. S5).

Our analysis suggests that SegB homologs of different lengths evolved via lineage-specific addition of domains of unknown function between two highly conserved ancestral domains. Since these seemingly diverse homologs were retained throughout evolution, one wonders about the ancestral, conserved function of all SegBs.

### The phylogenetic distribution of SegB correlates with ESCRT-III proteins and round cell morphology

To identify a biological context that might have favored emergence and maintenance of SegB as a component of the DNA segregation machinery, we looked for shared molecular and cellular features common to the *segB*-encoding Sulfolobaceae but not shared with nearby clades. In addition to being non-rod shaped thermophiles, all Sulfolobaceae contain a CdvB (ESCRT-III) homolog involved in cytokinesis. (Fig. 5). The ancestor of extant Sulfolobaceae was, therefore, likely also a non-rod shaped thermophile that employed ESCRT-III for division. The correlation between rounded cell morphology and the presence of a *segB* gene suggested to us that SegB may help break symmetry and establish axes of chromosome segregation and division in cells that lack an obvious long axis.

**Figure 5.**
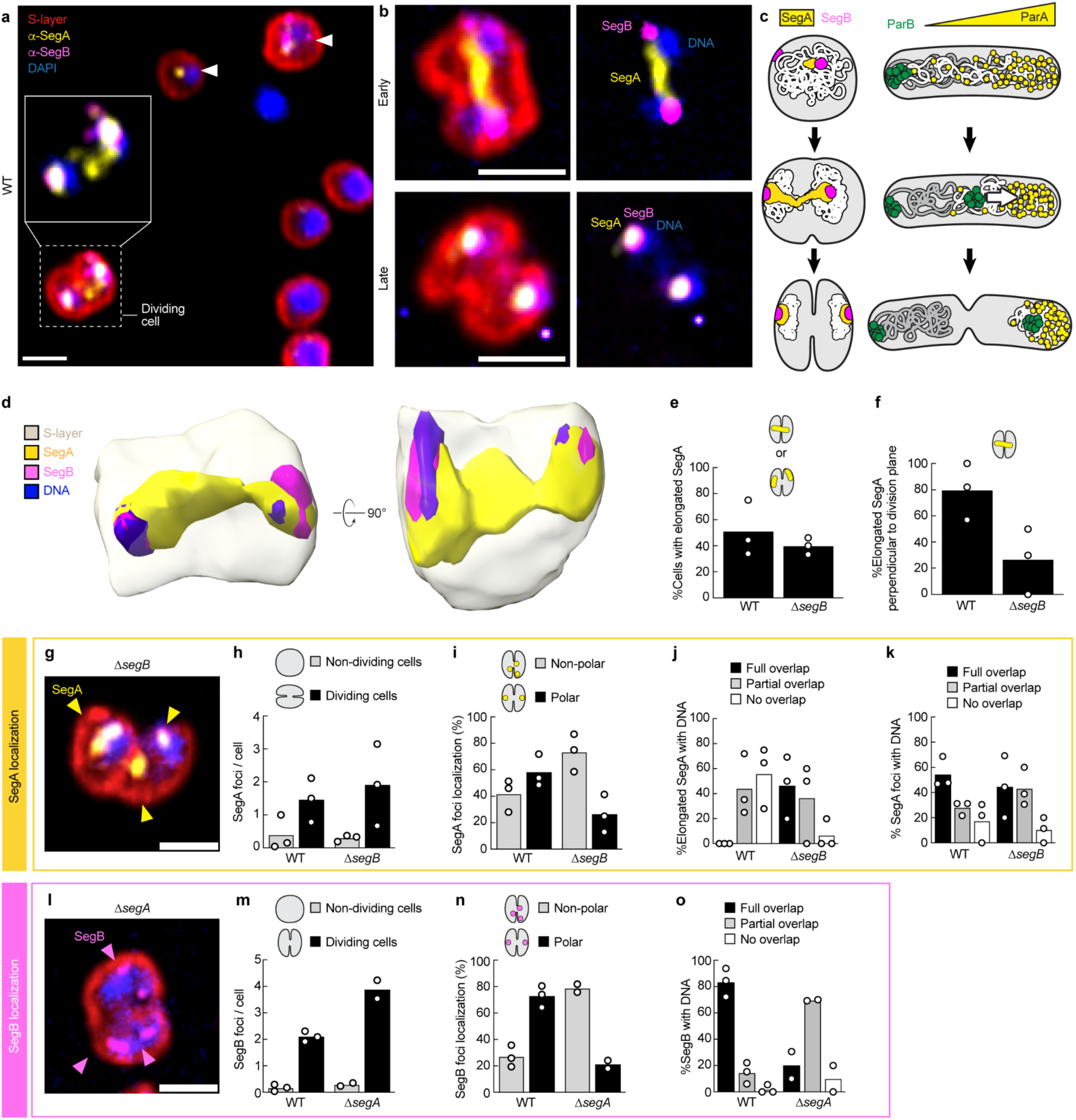
Subcellular localization and function of endogenous SegA and SegB in *Sulfolobus acidocaldarius*. **a.** Representative example of the subcellular localization of endogenous SegA (yellow) and SegB (magenta) in WT Saci obtained by immunofluorescence, soft-embedding in agarose, super-resolution imaging and deconvolution using specific antibodies. Chromosomal DNA was detected with DAPI (blue), and the S-layer was covalently labeled with an amine-reactive Alexa Fluor 647 (red). The inset shows an enlarged view of a dividing cell where the S-layer signal was removed for clarity. Arrowheads indicate non-dividing cells containing SegA and SegB foci. **b.** In earlier stages of cell division, SegA is frequently seen as an elongated structure stretching between two polar SegB foci and perpendicular to the division plane. In later stages, SegA typically presents as discrete foci that fully overlap with SegB and the DNA. **d.** Volumetric surface rendering of a *Saci* cell using ChimeraX software. The SegA elongated figure is clearly seen to span the two chromosomes and extend through the division plane. The gray cell surface was optimized using the *midsurf* script in ChimeraX. **e, f**. Occurence and orientation of elongated SegA structures in WT or in a strain of Saci where *segB* was deleted (Δ*segB*). **g.** Representative example of the subcellular localization of endogenous SegB in Δ*segB* and **(h-k)** corresponding quantification. **l.** Representative example of the subcellular localization of endogenous SegB in Δ*segA* and **(m-o)** corresponding quantification. A total of 161 (WT), 81 (Δ*segA*) and 109 (Δ*segB*) cells were analyzed; each dot represents the mean of a biological replicate.

### SegAB forms a bipolar figure that links segregating sister chromosomes and runs perpendicular to the plane of cell division

We investigated the subcellular localization of SegAB in *S. acidocaldarius* by generating antibodies against recombinant full-length SegA and SegB proteins from *Saci* and performing super-resolution immunofluorescence microscopy. Cells were soft-embedded in low-melt agarose to minimize mounting artifacts. Instantaneous Structured Illumination Microscopy (iSIM) revealed that SegB forms two discrete foci in 80% of all dividing cells that colocalize with segregated chromosomes, generally at or near the cell poles (Fig. 5a, b, d). SegA exhibited multiple localization patterns depending on whether the cell was early or late in the division process.

For quantitative analysis we separated images of dividing cells into Early and Late Stages based on chromosome and S-layer morphology. We scored cells as Early-Stage Division if they contained two compacted and clearly individuated chromosomes, but exhibited only shallow ingression of the cell envelope. We scored as Late-Stage Division cells containing two highly compacted, well-separated chromosomes with a deep cleavage furrow ingression. Remarkably, ∼50% of Early Stage Division cells contained a bipolar SegA/SegB figure composed of a dense, linear SegA structure capped at both ends by SegB foci. The SegA figure was invariably aligned perpendicular to the plane of cell division and did not correspond to an obvious underlying pattern of chromosomal DNA (Fig. 5a, c, d, Fig. S6). Volume rendering unambiguously shows that SegA structures span the whole cell and connect SegB foci from both cell poles (Fig. 5d). In 60% of Late Stage Division cells, SegA no longer formed a single, linear figure, but two separated foci, often colocalized with the separated SegB foci and chromosomal DNA at the cell poles (Fig. 5b). Less frequently, we observed intermediate patterns, usually with SegA and SegB forming separate puncta in Early-Stage Division cells, and overlapping/colocalized figures in Late-Stage Division cells (Fig. S6). These SegA and SegB localization patterns differ dramatically from those of ParA and ParB in rod-shaped bacteria (Fig. 5c) and point to fundamentally different mechanisms of chromosome segregation.

### SegA consolidates SegB foci while SegB orients the divisional SegA figure

To understand the specific functions of SegA and SegB we compared their localization patterns in wildtype and mutant cells using super-resolution immunofluorescence microscopy. In Δ*segA* cells, SegB no longer formed two discrete foci located at the cell poles during division, but instead formed an average of four foci that no longer localized to the cell poles and only partially overlapped chromosomal DNA (Fig. 5l, n). In Late Stage Division cells lacking *segB*, SegA still formed dense filament-like figures but they were less likely to be aligned perpendicular to the cell division plane (26% of Δ*segB* cells compared to 80% of WT cells, Fig. 5f). These elongated SegA figures were more strongly associated with chromosomal DNA (Fig. 6j). In Late Division Cells the number of SegA foci was unchanged in Δ*segB* cells, but these foci failed to localize to the cell poles. In both Δ*segA* and Δ*segB* deletion strains, chromosomal DNA appeared less frequently compacted than in WT cells, and even appeared fragmented in 27% and 15% of Δ*segA* and Δ*segB* cells, respectively, compared to 0% in WT cells (Fig. 1e). In cells where chromosomal compaction appeared normal, it was less frequently associated with the cell poles (only 57% and 69% in Δ*segA* and Δ*segB* cells, compared to 83% in WT cells, Fig. 1f). We conclude from these experiments that SegA and SegB drive chromosome segregation, but via a mechanism distinct from that of ParAB in rod-shaped bacteria.

## Discussion

Archaea are a phenomenally diverse but poorly understood domain of life. They are ubiquitous: thriving in oceans, briny pools, hot springs, soil, and animal (including human) microbiomes (Baker et al. 2020; Borrel et al. 2020; Duller and Moissl-Eichinger 2024; Hoegenauer et al. 2022). Archaea play an essential role in nitrogen and carbon cycling, and they produce most of earth’s atmospheric methane (Offre, Spang, and Schleper 2013; Garcia, Gribaldo, and Borrel 2022). In addition to their ecological significance, archaea also occupy a key position in the tree of life, as eukaryotes appear to result from at least one fusion event between a bacterium and an archaeon from the Asgard superphylum (Spang et al. 2015; Liu et al. 2021; Karki, Barth, and Aylward 2024; Spang et al. 2018; Zaremba-Niedzwiedzka et al. 2017; Krupovic, Dolja, and Koonin 2023; Eme et al. 2023; 2017).

Archaea share some chromosome segregation machinery with bacteria, especially deviant Walker-box ATPases of the ParA family. Interestingly, some archaeal species contain homologs of bacterial ParB proteins, while others appear to have innovated new regulatory proteins to control ATP hydrolysis by ParA homologs. Previous work in Sulfolobales identified two such archaea-specific regulators. The high copy number plasmid, pNOB8, from *Sulfolobus* sp. strain NOB8H2, for example, encodes a complete *parAB* operon, with ParA and ParB homologs that are ∼37% and ∼58% identical to bacterial counterparts. This operon also encodes an additional, archaea-specific protein, AspA, that mediates interaction between ParB and DNA (Schumacher et al. 2015). The SegAB operon represents a more extreme case, in which ParB has been replaced by either one (SegB) or two (SegB and SegC) archaea-specific factors that regulate SegA activity and promote chromosome segregation.

We find that SegB is not simply “ParB with different letters” but a SegA effector that facilitates an entirely different mechanism of chromosome segregation. Specifically, SegAB in *S. acidocaldarius* promotes bipolar chromosome partitioning by a mechanism distinct from that of ParAB-based systems in rod-shaped bacteria. For example, the localization patterns of SegA and SegB during division are not consistent with either the DNA relay model (Lim et al., 2014) or the observed distributions of ParA and ParB in rod-shaped bacteria. In *Caulobacter*, ParA is depleted from regions adjacent to ParB foci—especially from the region *between* segregating ParB/*parS* complexes. Late in the segregation process, migrating ParB foci appear to chase the ParA localization toward the distal pole. In contrast, SegA becomes highly *concentrated* in the region between the separating SegB/*segS* complexes, forming a dense, filament-like figure. The biophysical nature of this bipolar SegA/SegB figure is unclear, but it reminds us of early reports that SegB-induced ATP hydrolysis causes purified SegA to self-assemble into high order oligomers *in vitro* (Kalliomaa-Sanford et al., 2012). In late stages of division, as the cleavage furrow ingresses, the dense central SegA localization pattern splits in two and the separated halves retreat *toward* the segregated SegB foci.

Comparing SegAB and ParAB localization patterns reveals a fundamental difference in the functional relationships between the two sets of partner proteins. Rather than providing a target for diffusing DNA/protein segregation complexes, our data suggest that self-assembly of SegA either pushes chromosomes apart or prevents them from remixing before division is complete. We are reluctant to speculate on the molecular nature of the divisional SegA figure, but we note a couple of generic possibilities. SegA might form linear protein polymers similar to ParM in bacteria (Garner et al., 2007; Gayathri et al., 2012) or possibly a more amorphous phase condensate that is teased into an elongated pattern by association with trace amounts of DNA that linger between the segregating chromosomes. Interestingly, Kabli et al. (2025) recently reported that SegA forms DNA-associated foci in non-dividing *Sa. solfataricus* cells, a result consistent with our observations in *S. acidocaldarius*. These authors did not, however, report results for dividing cells, so it remains to be seen whether *Sa. solfataricus* SegA/SegB forms similar bipolar figures during division.

Regardless of the molecular nature of the bipolar SegA/SegB figures, their assembly must be relatively rapid (<180 seconds). We deduce this from the fact that they appear only during division—i.e. in cells with an obvious cleavage furrow. By live-cell, time-lapse imaging the division process, from furrow initiation to final cell separation, takes approximately 180 seconds, establishing this as an upper limit on the lifetime of the bipolar SegA/SegB figures (Charles-Orszag et al., 2021).

During division, the SegB/*segS* complex appears to provide critical spatial information required to align the entire bipolar SegA/SegB figure perpendicular to the plane of cell division. The SegB/*segS* complex also appears to promote detachment of SegA from the chromosome. We say this because loss of SegB does *not* prevent formation of an elongated divisional SegA figure, but randomizes its orientation and results in more spatial overlap between SegA and chromosomal DNA. Parham et al. (Parham et al., 2025) recently reported evidence of spatio-temporal coordination between the SegAB system and the ESCRT-III proteins that form the cytokinetic ring in Sulfolobales (Dobro et al., 2013; Pulschen et al., 2020; Samson et al., 2008; Tarrason Risa et al., 2020). We speculate that this coordination might underlie the SegB-dependent orientation of divisional SegA figures (Fig. 5f).

SegA also contributes—directly or indirectly—to chromosome compaction in *S. acidocaldarius*. In Sulfolobales, chromosome compaction occurs locally via the small nucleoid associated proteins Alba, Cren7, and Sso7d (Cajili and Prieto, 2022; Laursen et al., 2021); larger chromosome compartments are organized by the SMC-like protein coalescin (Takemata et al., 2019; Takemata and Bell, 2020). In addition, in contrast to the single origin of replication in bacterial chromosomes, the chromosome in Sulfolobales contains three origins of replication (Lundgren et al., 2004) that are believed to associate in the cell, in part due to the action of coalescin (Takemata et al., 2019). It is possible that SegA may interact with these proteins. In any case, our observations are consistent with other recent studies (Parham et al., 2025) and suggest that chromosomal ParA-like proteins have evolved additional functions in archaea. The DNA compaction activity in *Saci* SegA is important in that it seems to promote formation of tightly focused SegB-DNA clusters, possibly to ensure segregation of all three origins of replication at once. Loss of SegA causes the two discrete foci of SegB observed in wildtype cells to fragment into multiple spots that fail to find the cell poles. Our data, as well as other recent work (Kabli et al., 2025), reveals that Saci SegB binds both a centromeric sequence *segS*, and more distal sites on the chromosome. SegB cluster formation may be mediated by SegB disordered regions in *Saci* that are known drivers of phase separation (Borcherds et al., 2021), or by oligomerization, as residues proposed to promote oligomerization in *Sso* SegB are conserved and solvent-exposed in *Saci* SegB. In other species, phase separation may be promoted by the extensive N-terminal insertions. In any case, the concentration of SegB into a single focus on DNA requires SegA.

Finally, loss of either SegA or SegB causes a growth defect in both *Saci* (this study) and *Sso* (Kabli et al., 2025), but the genes encoding these proteins are not essential in either organism. The same is true for *parAB* operons in many bacterial species, such as *Deinococcus radiodurans*, *Vibrio cholerae*, *Bacillus subtilis*, *Pseudomonas aeruginosa*, and *Mycobacterium smegmatis* (Charaka and Misra, 2012; Donovan et al., 2010; Du et al., 2016; Jecz et al., 2015; Kadoya et al., 2011; Lee and Grossman, 2006; Lewis et al., 2002; Minnen et al., 2011; Santi and McKinney, 2015). Additional, robust but cryptic DNA-segregation mechanisms must exist in both bacteria and archaea. These mechanisms could involve either proteins specially adapted to segregation or non-specific biophysical driving forces such as entropy or DNA looping (Goloborodko et al., 2016; Harju et al., 2024; Jun and Wright, 2010). Regardless, it appears that the ParAB and SegAB systems have evolved to employ similar ATPases but different biophysical mechanisms to accelerate chromosome segregation, make it more reliable, and/or coordinate it with cell division. Our work sheds light on the diversity of mechanisms for ParA-based chromosome segregation, and may possibly foreshadow additional species-specific differences—in other archaea but also in non-rod-shaped bacteria where ParABS mechanisms are far from being completely understood.

## Methods

### Cell culture and preparation of conditioned media

All *Sulfolobus acidocaldarius* strains were grown aerobically with agitation at 75°C in Brock’s media (Brock et al., 1972) supplemented with 0.1% tryptone, 0.2% dextrin and 10 ug.mL^-1^ uracil (Sigma-Aldrich). Conditioned Brock’s media was prepared by centrifugation of 50 mL of an exponentially growing culture (OD_600nm_ ∼0.2-0.3) at 4,000 xg and room temperature for 15 min followed by filtration through a 0.22 µm filter to remove cells and cell debris, and was stored at 4°C for up to a month. Cell growth of WT and deletion strains was assessed by monitoring optical density at 600 nm.

### Construction of in-frame markerless deletion mutants in *S. acidocaldarius*

Knock-out strains were generated via the pop-in pop-out method described in (Wagner et al., 2012). For the *segA* knockout, 557 bp upstream and 676 bp downstream of *segA* were PCR amplified using primers 13461/13462 and 13463/13464, connected via overlap PCR using primers 13461/13464, and cloned between the NotI and NdeI sites of pSVA407, resulting in plasmid pSVA6463. For the *segB* knockout, 606 bp upstream and 552 bp downstream of *segB* were PCR amplified using primers 13457/13458 and 13459/13460, connected via overlap PCR using primers 13457/13460, and cloned between the NdeI and BamHI sites of pSVA407 resulting in plasmid pSVA6462. Plasmid transformation in the parental strain MW001 (referred to as WT throughout the manuscript), initial selection via blue/white screening, secondary selection, and mutant screening were performed as previously described. The resulting deletion mutants MW1264 (Δ*segA*) and MW1263 (ΔsegB) were confirmed by sequencing.

### Cloning and expression of recombinant SegA and SegB from *S. acidocaldarius*

All the sequences from *S. acidocaldarius* were codon optimized for expression in *E. coli* and ordered as synthetic DNA fragments (Integrated DNA Technologies). The fragment for His_6_-KCK-SegA was ligated between the NdeI and NcoI sites of the pET-22b(+) (Novagen) resulting in plasmid pACO29. The fragment for SegA-His_6_ was ligated between the NdeI and XhoI sites of the pET-22b(+) as in (Kalliomaa-Sanford et al. 2012) resulting in plasmid pACO50. The gene fragments for SegB and SegB_S56C_ were ligated between the SapI and XhoI sites of a N-His-ySUMO expression vector (pTB146) resulting in plasmids pACO3 and pACO24, respectively. All constructs were expressed in *E. coli* Rosetta2 cells. Expression was induced with 1 mM Isopropyl β-D-1-thiogalactopyranoside (IPTG) when cultures reached OD_600nm_ ∼2-3 and carried out overnight at 16°C. Cells were lysed in a Microfluidizer LM10 (Microfluidics) in respective His-tag binding buffers containing 1 mM phenylmethylsulfonyl fluoride (PMSF) and a protease inhibitor cocktail (Roche). Lysates were clarified by ultracentrifugation at 35,000 x*g* and the supernatants injected onto a HisTrap excel column (Cytiva). His-tagged proteins were eluted with a gradient of 30 mM to 400 mM imidazole. Protein-containing fractions were pooled and dialyzed overnight against storage buffer. For SUMO-tagged proteins, the tag was removed by the addition of yeast His_6_-Ulp1 SUMO protease purified in-house overnight at 4°C during dialysing into His-tag binding buffer. Both the tag and the protease were removed from the protein mixture by passage onto a Ni-NTA agarose resin (Qiagen) by gravity followed by gel filtration HiPrep 26/60 Sephacryl S100 HR column (Cytiva). <u>For His_6_-KCK-</u> <u>SegA:</u> His-tag binding buffer was 20% sucrose, 25 mM HEPES-KOH pH 7.4, 500 mM KCl, 20 mM imidazole, 2 mM MgCl_2_, 5 mM β-Mercaptoethanol (βME), 500 mM Mg-ATP; elution buffer was 20% sucrose, 25 mM HEPES-KOH pH 7.4, 500 mM KCl, 400 mM imidazole, 2 mM MgCl_2_, 5 mM βME, 100 mM Mg-ATP; storage buffer was 20% sucrose, 25 mM HEPES-KOH pH 7.4, 500 mM KCl, 2 mM MgCl_2_, 1 mM tris(2-carboxyethyl)phosphine (TCEP), 100 mM Mg-ATP. <u>For SegA-His_6_:</u> His-tag binding buffer was 10% glycerol, 20 mM Tris-HCl pH 7.4, 300 mM NaCl, 20 mM imidazole, 2 mM MgCl_2_, 500 mM Na-ATP, 5 mM βME; elution buffer contained 400 mM imidazole; storage buffer was 10% glycerol, 20 mM Tris-HCl pH 7.4, 2 mM MgCl_2_, 500 mM Na-ATP, 2 mM TCEP. <u>For all SegB constructs:</u> binding buffer was 30 mM Tris-HCl pH 7.4, 250 mM NaCl, 30 mM imidazole, 5 mM βME; elution buffer was identical but contained 300 mM imidazole; dialysis/tag cleavage buffer contained 30 mM imidazole and 2 mM TCEP; storage buffer 20% glycerol, 30 mM Tris-HCl pH 7.4, 250 mM NaCl, 2 mM TCEP (note: all pH values provided correspond to pH at 4°C).

### Electrophoretic mobility shift assays (EMSA)

For EMSAs, proteins were buffer exchanged in DNA binding buffer (10 mM Tris-HCl pH 7.4, 50 mM KCl, 5 mM MgCl_2_, 2.5% glycerol, 0.01% NP-40). Synthetic DNA fragments (Integrated DNA Technologies) stocks were resuspended at 1 mM in nuclease-free water. For DNA fragments larger than 50 bp, reaction mixes were subjected to polyacrylamide gel electrophoresis in NuPAGE™ Bis-Tris Mini Protein Gels, 4–12% (Invitrogen) in Tris-Glycine (TG) 1X buffer at 4°C. For DNA fragments smaller than 50 bp, reaction mixes were subjected to 3% agarose gel electrophoresis in Tris-borate-EDTA (TBE) 0.5X buffer at room temperature. DNA was stained with 1X SYBR Gold and destained in either TG 1X or TBE 0.5X. Gels were imaged in a ChemiDoc imager (BioRad).

### DNA curtain assays

DNA curtains were established as previously described (Gallardo et al., 2015, Keenen et al., 2021). Briefly, chrome features (or barriers) were nanofabricated on microscope slides which were then converted to microfluidic flow-cells. Liposomes were used to create a fluid lipid bilayer around chrome barriers on the surface of a flow-cell. Biotinylated DNA from bacteriophage lambda (λ-DNA) was tethered to individual biotinylated lipids via a multivalent streptavidin interaction. Buffer flow was then used to align and extend tethered λ-DNA over chrome barriers to create visualizable DNA curtains stained with the intercalating dye YOYO-1. A solvent-accessible cysteine was introduced in SegB by point mutation (S56C), allowing cysteine labeling with Alexa Fluor 569 C5 maleimide according to the manufacturer’s instructions (Thermo Fisher Scientific). SegB_S56C_-AF568 (50% labeled) was injected into the flow-cell at 1µM in imaging buffer containing 2 0mM Tris-HCl (pH 7.5), 50mM NaCl, 0.5mg/mL BSA, 0.5% Pluronic F-127, 1 mM MgCl_2_, 1mM DTT, and 10 pM YOYO-1. SegB_S56C_-AF568 binding to DNA was imaged at 0.1 Hz with a buffer flow rate of at least 0.5mL/min to maintain DNA extension at chrome barriers. DNA curtains were imaged using total internal reflection fluorescence (TIRF) microscopy, and images were captured with an EMCCD camera run on Micro-Manager and Fiji. Binding distributions of SegB_S56C_-AF568 were measured in Python using analysis described previously (Ichino et al., 2021, Strauskulage et al., 2023). Briefly, fluorescent channels were aligned and individual λ-DNA molecules were identified in an image based on YOYO-1 signal. These coordinates were used to identify SegB_S56C_-AF568 binding events on DNA and two-dimensional gaussians were fitted to determine the sub-pixel position of each SegB_S56C_-AF568 molecule. The relative base pair position of each SegB_S56C_-AF568 binding event was calculated based on its distance from the free-end of the DNA using the estimated extension of tethered, flow-extended λ-DNA. The mapped SegB_S56C_-AF568 binding distribution along λ-DNA was plotted alongside the distribution of “AGNGT” binding sites identified using SeqIO from Biopython.

### Protein crystallization and structure determination

SegB was used at a concentration of 11-18 mg/mL. Sitting drop vapor diffusion trays were set up and incubated at room temperature, containing 0.4 µL drops of protein and reservoir solution in a 1:1 ratio. Crystals formed after 35 days in 100 mM NH_4_Cl and 17-19% PEG 3350. Phases were determined by soaking crystals for 2 hours in a solution consisting of mother liquor and 2 mM K_2_PtI_6_. Data was collected at ALS 8.3.1 on a Pilatus3 S 6M detector. Data indexing was completed using XDS and AIMLESS in the CCP4 suite. Structure determination and refinement were done in the Phenix programming suite. Visualization of SegB structure for figure preparation was done in ChimeraX.

### Circular dichroism spectroscopy

Circular dichroism spectroscopy was performed in a Jasco J715 spectrometer with 10 μM SegB in 10 mM potassium phosphate buffer pH 7.4. Buffer alone was used as a blank. SegB secondary structure at 30°C was determined by measuring ellipticity from 200 nm to 280 nm. SegB thermostability was determined by measuring ellipticity at 222 nm while varying the temperature from 30°C to 110°C. SegB secondary structure was determined again by lowering the temperature back to 30°C and measuring ellipticity from 200 nm to 280 nm.

### Bioinformatic analyses

Phylogenetic analyses were done in Geneious Prime (Biomatters Ltd.). A homology search was performed in BLASTP against the nucleotide collection database (nr/nt) using *Sa. solfataricus* SegB as query. Hits were manually filtered for duplicates or poorly annotated sequences, yielding a final list of 72 hits. These sequences were aligned with the Geneious Alignment tool, and a SegB homologs tree was built with the Geneious Tree Builder using a Neighbor-Joining method to account for potential differences in evolution rates between lineages. To map SegB homologs onto the natural phylogenetic tree of archaea, fifty-seven 16s rRNA sequences were manually retrieved from the National Center for Biotechnology Information (NCBI) database corresponding to SegB-encoding and SegB-non-encoding Euryarchaea and Thermoproteota archaeal species, and included the 16s rRNA sequence of *Escherichia coli* as an outgroup. Sequences were aligned with the Geneious Alignment tool and the tree was built with the Geneious Tree Builder using Unweighted Pair-Group Method with Arithmetic Averaging (UPGMA). Known or predicted presence or absence of ParA-like, ParB, and CdvB proteins in these 57 species was assigned manually using available literature and NCBI BLASTP.

### Antibodies

Polyclonal antibodies were raised against full-length recombinant tag-less SegB in rabbit and His_6_-KCK-SegA in guinea pig (Covance). Anti-SegB antibodies were further affinity-purified against full-length SegB crosslinked to AminoLink Coupling Resin (ThermoFisher) according to the manufacturer’s instructions, and used at ∼0.6 µg.mL^-^_1_ (∼1:1,000) for immunostaining. Anti-SegA was cleaned up by ultracentrifugation at 65,000 xg for 30 min and the supernatant used at 1:1,000. Secondary goat-anti-rabbit AlexaFluor 488 antibodies and goat-anti-guinea pig AlexaFluor 568 antibodies (ThermoFisher) were used at 1:1,1000. All antibodies were diluted in 3% bovine serum albumin in phosphate buffered saline containing 0.05% Tween-20 (PBST-3% BSA).

### Immunostaining and soft embedding in low melting point agarose

15 mL of exponentially growing cells (OD_600nm_ ∼0.2-0.3) were collected by centrifugation at 4,000 xg for 15 min at room temperature and the supernatant was spared. Cells were then washed once in PBS 1X, resuspended in 225 µL PBS 1X and added to a 1.5 mL microcentrifuge tube. The S-layer was labeled by adding 25 µL of AlexaFluor 647-NHS-ester (ThermoFisher) freshly reconstituted at 0.5 mg.mL-1 in anhydrous dimethyl sulfoxide (DMSO). Cells were incubated in the dark at room temperature with gentle agitation for 20 minutes, after which they were centrifuged at 4,000 xg for 5 min, washed once with PBS 1X and once with non-pH-adjusted minimal Brock media, and added back to the original culture supernatant in a culture flask and allowed to grow for 1,5h. Cells were then collected by centrifugation at 4,000 xg for 15 min at 40°C, washed with prewarmed citrate buffer pH 3, and fixed stepwise in increasing concentration of ice-cold ethanol according to (Cezanne et al., 2023). Fixed cells were collected by centrifugation at 4,000 xg for 15 min at room temperature, washed twice in PBS 1X, resuspended in 250 µL permeabilization buffer (GTET buffer: 50 mM glucose, 20 mM Tris pH 7.5, 10 mM EDTA, 0.2% Tween-20) and incubated in the dark at room temperature for 20 min with gentle agitation. Permeabilized cells were collected by centrifugation at 4,000 xg for 5 min at room temperature, rinsed three times in PBS 1X, and blocked in 250 µL PBST-3% BSA overnight at 4°C. Anti-SegA and anti-SegB antibodies were added and incubated for 1h at room temperature with gentle agitation. Cells were washed four times in PBS 1X, resuspended in 250 µL PBST-3% BSA containing fluorescently labeled secondary antibodies and 10 µg.mL-1 4’,6-diamidino-2-phenylindole (DAPI, ThermoFisher), incubated for 1h at room temperature with gentle agitation, washed four times in PBS 1X and resuspended in 1 mL PBS 1X. 1-10 µL cells were added to 90 µL PBS 1X in wells of a 96-well glass bottom plate (SensoPlate, Greiner BioOne) that was previously plasma treated, coated with 0.01% poly-L-lysine (ThermoFisher) for 30 min at room temperature, washed with MilliQ water and air-dried. The cell-containing plate was centrifuged at 1,000 xg for 2 minutes at room temperature. Cells were soft-embedded by slowly adding 100-150 µL of a solution of 0.5% low melting point agarose (UltraPure LMP agarose, Invitrogen) in 0.4% sucrose and 0.1M NaCl at 40°C after carefully discarding the supernatant. LMP agarose was allowed to set at least 30 min at room temperature prior to imaging.

### Super-resolution fluorescence imaging

Immunostained cells were imaged on a iSIM (Visitech) equipped with 405, 488, 561 and 647 nm lasers, a Nikon 100X 1.45NA PlanApo objective and a 1.5X optovar lens, and a Hamamatsu Quest CMOS camera. Z-stacks were acquired with 0.125 µm steps. Raw images were deconvolved with Microvolution in the Micro-Manager software (Edelstein et al., 2010).

### 3D volumetric surface rendering

Even though the S-layer can be approximated to a 2D surface with virtually no depth at the cellular scale, it appears as a ∼200 nm-thick layer due to optical resolution limitations. To facilitate interpretation of SegA and SegB subcellular localizations, we developed a custom code in Python, *midsurf*, to recalculate the theoretical position of the S-layer in ChimeraX (Goddard et al., 2018). This code moves surface vertices along a volume gradient until they reach a maximum to find the center S-layer surface in a roughly spherical shell (https://rbvi.github.io/chimerax-recipes/middle_surface/midsurf.html).

### High-temperature live-cell imaging

High-temperature live-cell imaging was done as in (Charles-Orszag et al., 2021). Briefly, a DeltaT cell micro-environment control system (Bioptechs) was modified by the manufacturer to allow it to reach temperatures up to 80°C. It was used on a Nikon Ti-E inverted microscope equipped with a motorized stage. In order to enhance cell attachment, the coverslips of the imaging chambers were modified with micropatches of 0.8% gelrite (Gelzan, Sigma-Aldrich). The microscope objective was prewarmed to 65°C prior to imaging and kept at this temperature throughout the experiment. Upon imaging, 500 µL of cells from an exponentially growing culture (OD_600nm_ ∼0.2-0.3) were added to an imaging chamber (final volume 2.5 mL) and allowed to settle for 15 min at 75°C. The chamber controller was set to 78°C to maintain cells and media between 74.5 and 75.6°C throughout the experiment. A heated glass lid was used to prevent condensation and evaporation. Differential interference contrast (DIC) microscopy was performed with a green light-emitting diode illuminator (B180-RGB; ScopeLED) through a Nikon 0.72NA CLWD air condenser, a Nikon PlanApo VC 100X 1.4NA oil objective, Cargille 1809 immersion oil (refractive index ∼1.515 at 65°C at 589nm), with an additional 1.5X optovar lens, and a PointGrey CMOS camera (CM3-U3-50S5M-CS). Cells were imaged with a 100-ms exposure every 5 seconds. All microscopy hardware was controlled with Micro-Manager software (Edelstein et al., 2010).

## Data analysis

Image analysis and quantifications were manually made in Fiji (ImageJ (Schindelin et al., 2012)). Data was analyzed and graphs were prepared in Prism (GraphPad). Figures and illustrations were prepared in Adobe Illustrator.

**Table 1.** Plasmids used in this study.

| Plasmid | Description | Source |
| --- | --- | --- |
| pACO3 | *Saci segB* gene (*saci0203*) codon optimized for expression in *E. coli*, ligated between SapI and XhoI sites of N-His-ySUMO expression vector (pTB146) | This study |
| pACO24 | *Saci segB* gene (*saci0203*) containing a S56C mutation, codon optimized for expression in *E. coli*, ligated between SapI and XhoI sites of N-His-ySUMO expression vector (pTB146) | This study |
| pACO29 | *Saci segA* gene (*saci0204*) codon optimized for expression in *E. coli*, with in frame N-terminal His<sub>6</sub> and KCK tags, ligated between the NdeI and NcoI sites of the pET-22b(+) (Novagen) | This study |
| pACO50 | *Saci segA* gene (*saci0204*) codon optimized for expression in *E. coli*, with in frame C-terminal His<sub>6</sub>, | This study |
|  | ligated between the NdeI and XhoI sites of pET-22b(+)<br>(Novagen) |  |
| pSVA6462 | *segB* deletion plasmid in *Saci* | This study |
| pSVA6463 | *segA* deletion plasmid in *Saci* | This study |

**Table 2.**
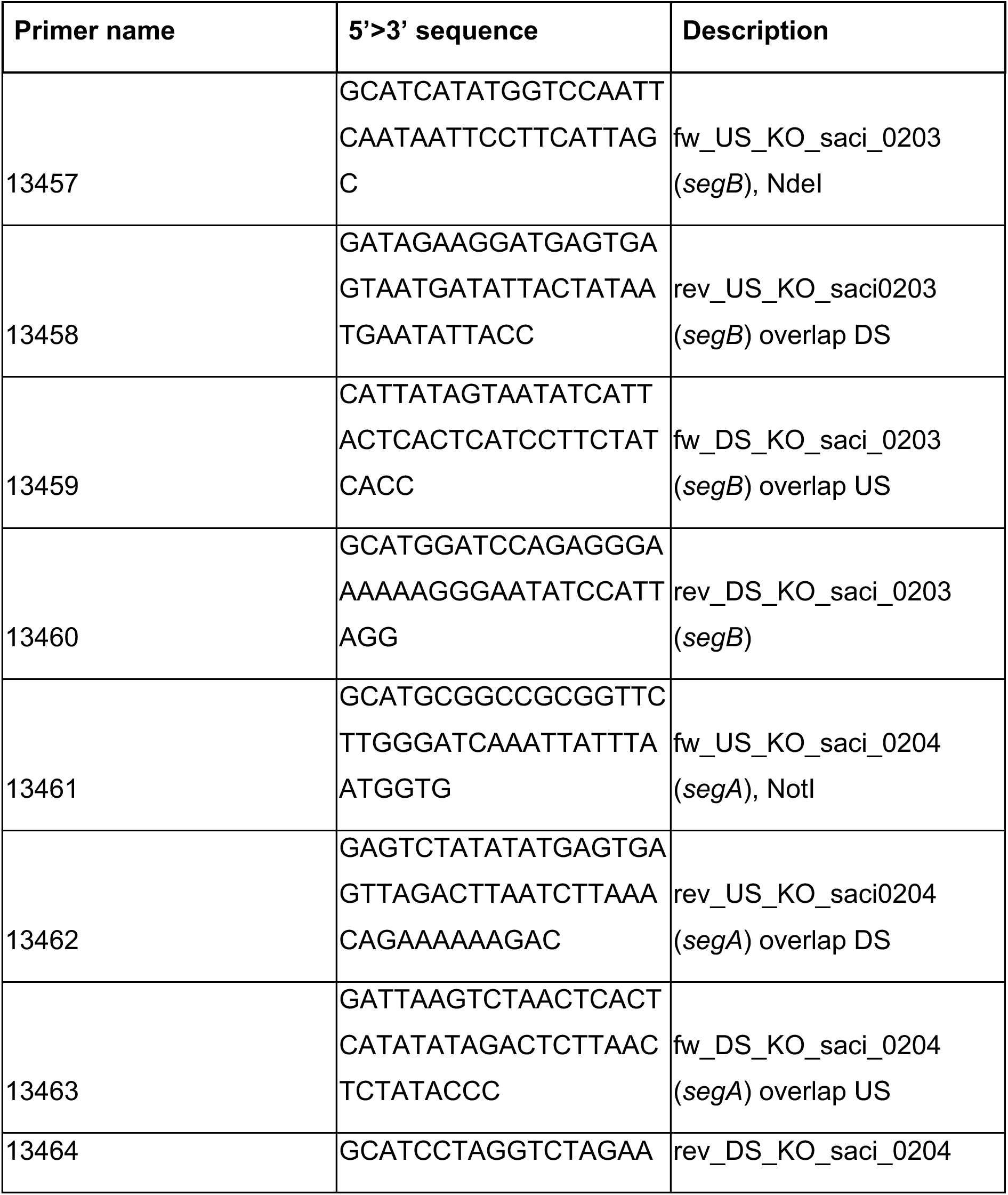
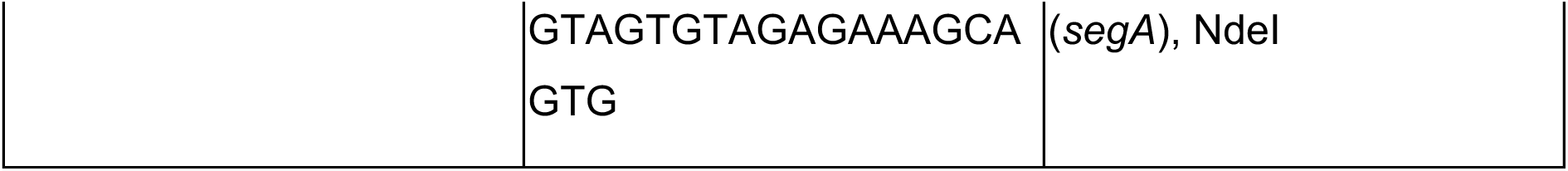
Primers used in this study.

**Table 3.** X-ray diffraction data and refinement statistics of SegB.

|  |  |
| --- | --- |
| **Wavelength** |  |
| **Resolution range** | 58.32 - 1.85 (1.9 - 1.85) |
| **Space group** | P 21 21 21 |
| **Unit cell** | 44.944 54.296 116.634 90 90 90 |
| **Total reflections** | 162629 |
| **Unique reflections** | 25076 (1771) |
| **Multiplicity** | 6.5 |
| **Completeness (%)** | 99.81 (99.94) |
| **Mean I/sigma(I)** | 21.6 |
| **Wilson B-factor** | 31.13 |
| **R-merge** | 0.042 |
| **R-meas** | 0.05 |
| **R-pim** | 0.026 |
| **CC1/2** | 1 |
| **Reflections used in refinement** | 25076 (1771) |
| **Reflections used for R-free** | 1999 (142) |
| **R-work** | 0.3193 (0.3680) |
| **R-free** | 0.3445 (0.4338) |
| **Number of non-hydrogen atoms** | 2170 |
| **macromolecules** | 1994 |
| **ligands** | 0 |
| **solvent** | 176 |
| **Protein residues** | 244 |
| **Nucleic acid bases** | n/a |
| **RMS(bonds)** | 0.008 |
| **RMS(angles)** | 1.04 |
| **Ramachandran favored (%)** | 96.14 |
| **Ramachandran allowed (%)** | 3.86 |
| **Ramachandran outliers (%)** | 0 |
| **Rotamer outliers (%)** | 0.89 |
| **Clashscore** | 67.51 |
| **Average B-factor** | 32.35 |
| **macromolecules** | 31.97 |
| **ligands** | n/a |
| **solvent** | 36.64 |

## Data availability

The atomic coordinates and structure factors have been deposited in the Protein Data Bank (https://www.rcsb.org/pdb) with PDB ID code 9OQF.

## Acknowledgments

This work was supported by the National Institute of General Medical Sciences of the National Institutes of Health (R35-GM118119 to RDM), by the Howard Hughes Medical Institute Investigator Program (RDM), and by a Momentum grant (VW Foundation grant number 94993 to SVA). ACO has received support by the Simons Foundation as a Life Sciences Research Foundation fellow. NH was supported by a UCSF Pulmonary Division Training Grant (2T32HL007185), the Burroughs Wellcome Fund Postdoctoral Enrichment Program (1019894), and a University of California President’s Postdoctoral Fellowship Program. We thank the X-ray Crystallography Facility at the University of California, San Francisco. We appreciate the use of Beamline 8.3.1 at the Advanced Light Source, which is supported by the University of California Office of the President MRPI program, Plexxikon Inc., and the Integrated Diffraction Analysis Technologies program of the US Department of Energy Office of Biological and Environmental Research. The Advanced Light Source (Berkeley, CA) is a national user facility operated by Lawrence Berkeley National Laboratory on behalf of the US Department of Energy under contract number DE-AC02-05CH11231, Office of Basic Energy Sciences. Molecular graphics and analyses performed with UCSF ChimeraX, developed by the Resource for Biocomputing, Visualization, and Informatics at the University of California, San Francisco, with support from National Institutes of Health R01-GM129325 and the Office of Cyber Infrastructure and Computational Biology, National Institute of Allergy and Infectious Diseases. The authors are particularly grateful to Jose De La Torre, Simonetta Gribaldo, Buzz Baum and Joe Parham, Daniela Barilla, Ethan Garner, Alex Bisson, Mecky Pohlschröder, Mullins lab members Justin Salat, Terri Lee, Jiongyi Tan, Natalie Petek, Kevin Nie, Wilson Nieves, Jessica Zhang, former students of the Physiology Course at the Marine Biological Laboratory, and former participants of the Archaeal Cell Biology Workshop, for years of fruitful discussions and technical help.

## Contributions

A.C.O.: conceptualization, methodology, investigation, formal analysis, validation, visualization, figure preparation, writing original draft preparation, and review and editing. S.L.: methodology, formal analysis, validation, visualization, figure preparation, writing original draft preparation, and review and editing. N.H., L.S., A.B., B.W., M.V.W., G.A., A.F., T.G. & J.R.: conceptualization, methodology, investigation, formal analysis, validation, visualization, and editing. S. R., O.R., S.V.A. & R.D.M.: funding acquisition, supervision, methodology, investigation, formal analysis, writing original draft preparation, and review and editing. All authors contributed to the article and approved the submitted version.

## Supplementary material

**Figure S1.**
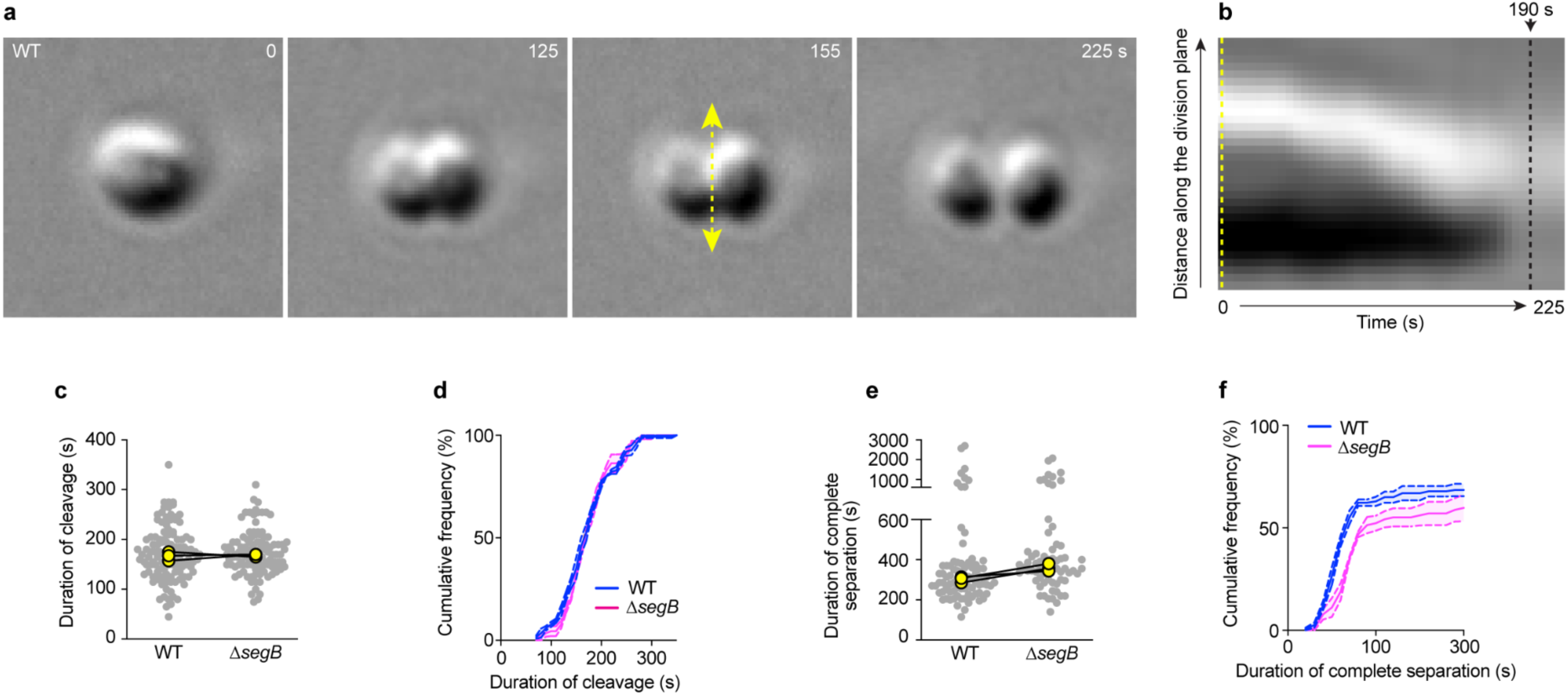
Live-cell imaging of cell division. **a.** Representative example of a dividing WT *S. acidocaldarius* cell imaged at 76°C using DIC. **b.** Kymograph showing the analysis of the duration of cleavage of the cytokinetic furrow. **c,d.** Duration of cleavage (time from the first invagination of the membrane to the point at which the cleavage is complete) and the time it takes the two daughters to fully separate for wildtype and *segB*-deletion mutant. Cells that were never observed to separate were excluded. Small gray dots represent each cell, while the large yellow dots are the medians of 3 independent biological replicates. **e,f.** Cumulative frequency distributions of all cells, including those that never separated. segB-deletion mutant cells exhibited a normal cleavage time, but then the daughters tended to stay connected longer. A slightly larger fraction of the *segB*-deletion cells never separated before the movies ended. Dark lines are the mean curves and light shading is the standard error of the mean from 3 biological replicates.

**Figure S2.**
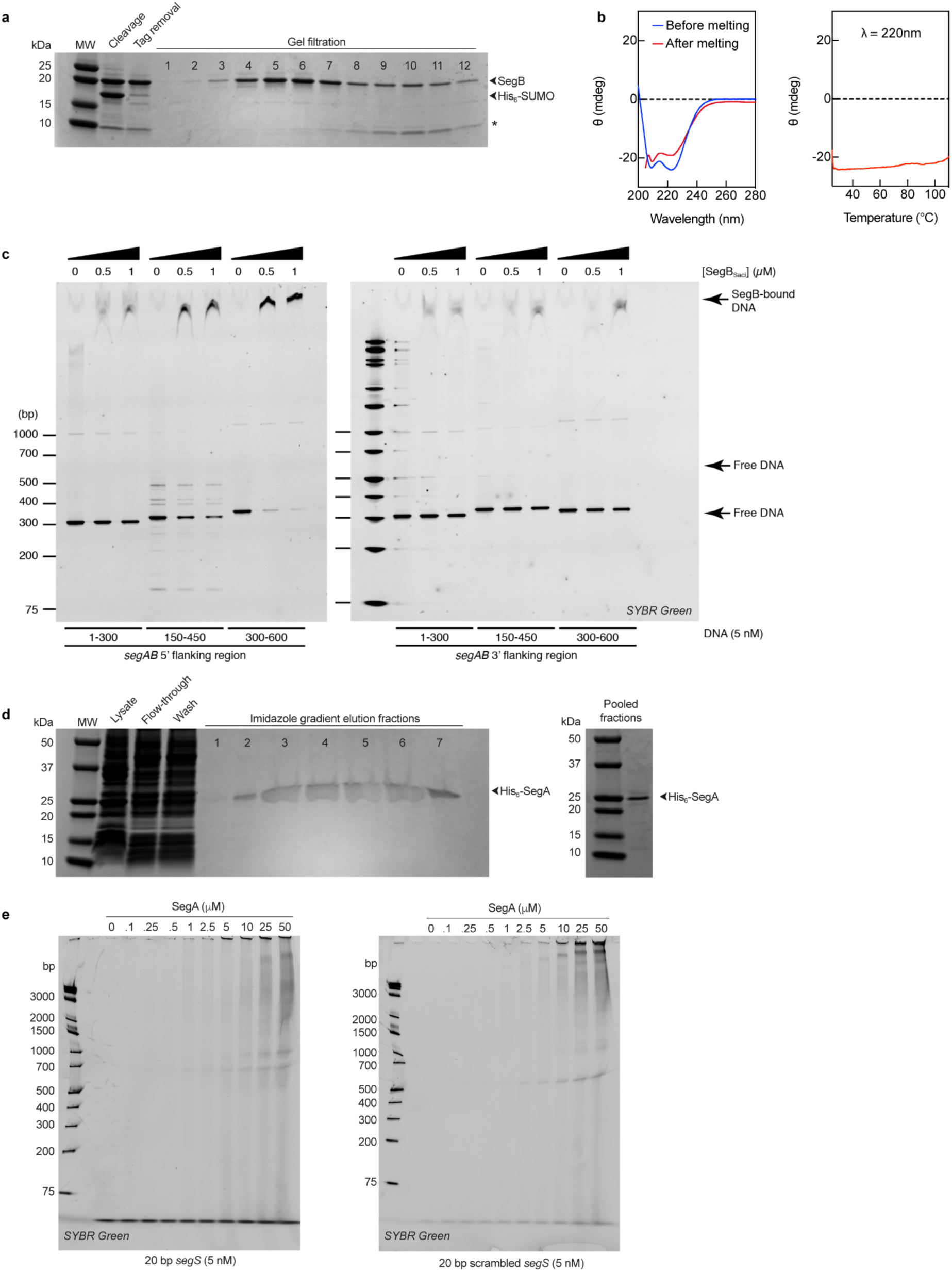
Protein purification, secondary structure determination of SegB, and DNA binding assays. **a.** Purification of SegB by gel filtration on a HiPrep 26/60 Sephacryl S100 HR column after nickel-affinity chromatography and enzymatic tag cleavage. **b.** Circular dichroism spectroscopy measurements of SegB ellipticity from 220 nm to 280 nm before and after melting at 110°C, showing that SegB mostly consists of ɑ-helices, and melting curve at 222 nm, showing that SegB remains folded at temperatures up to 110°C. **c.** Electrophoretic mobility shift assays (EMSAs) of SegB binding to DNA. Gels were stained for DNA using SYBR Green. (*Left*) SegB protein binds readily to the DNA matching the sequence of the region flanking the 5’ of the *segAB* operon, which contains the *segS* sequence we identified. At the higher SegB concentrations, the amount of free DNA is reduced (right columns). (*Right*) SegB also binds nonspecifically to other DNA sequences, but not with as high an affinity, as indicated by the reduced shifted DNA. **d.** Purification of recombinant SegA-His_6_. (*left*) Gel of the fractions eluted from a nickel-affinity column with imidazole. (*right*) Gel of pooled fractions showing purified SegA-His_6_. **e.** EMSAs of SegA binding to DNA either with (*left*) or without (*right*) the specific *segS* sequence. Gels were stained for DNA using SYBR Green. Increasing SegA concentrations as shown across the top bind DNA in a monotonic manner, indicated by the DNA shift increasing from left to right. SegA binds equally to *segS* and scrambled *segS* sequences, indicating nonspecific DNA binding.

**Figure S3.**
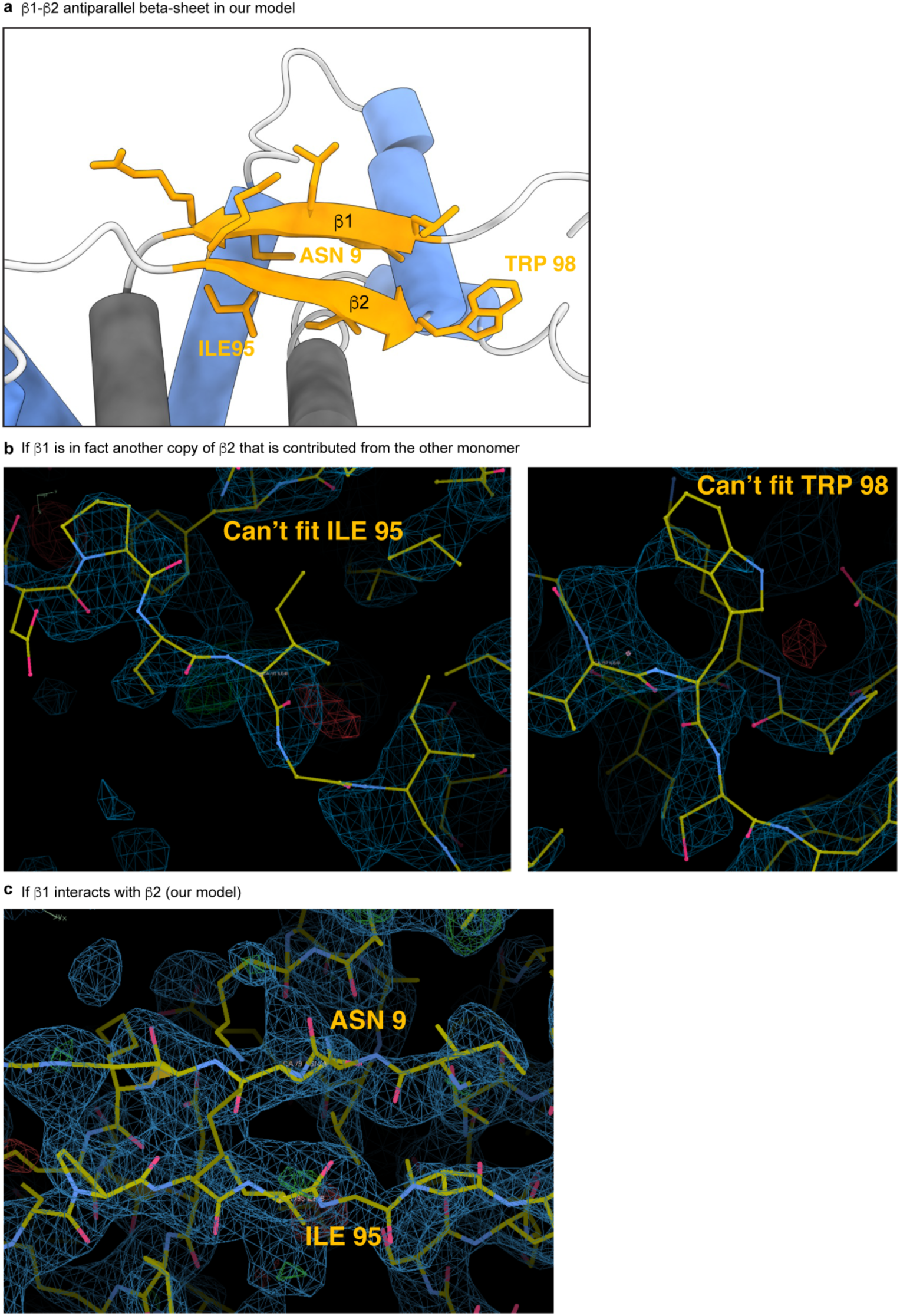
Details of the β-sheet region in SegB monomer. **a.** Detailed region of our crystal structure, which has two internal β strands stacking. The previously published structure of SegB from Sso had mirror β strand copies from two different monomers stacking to create the dimer interface. **b.** To test whether the stacking β strands are possibly from two different monomers, as seen in the previous structure, we examined the electron density of β1 in detail. If the β strands were mirror copies from different monomers, we would expect to see electron density from isoleucine 95 and tryptophan 98 from the same β sheet at the stacking region. But neither amino acid fits the observed electron density from our crystal structure data. **c.** Only asparagine from β1 and isoleucine 95 from β2 fit the electron density in the β stacking region, a sequence that corroborates our structural model that the two β strands are indeed internal to one monomer.

**Figure S4.**
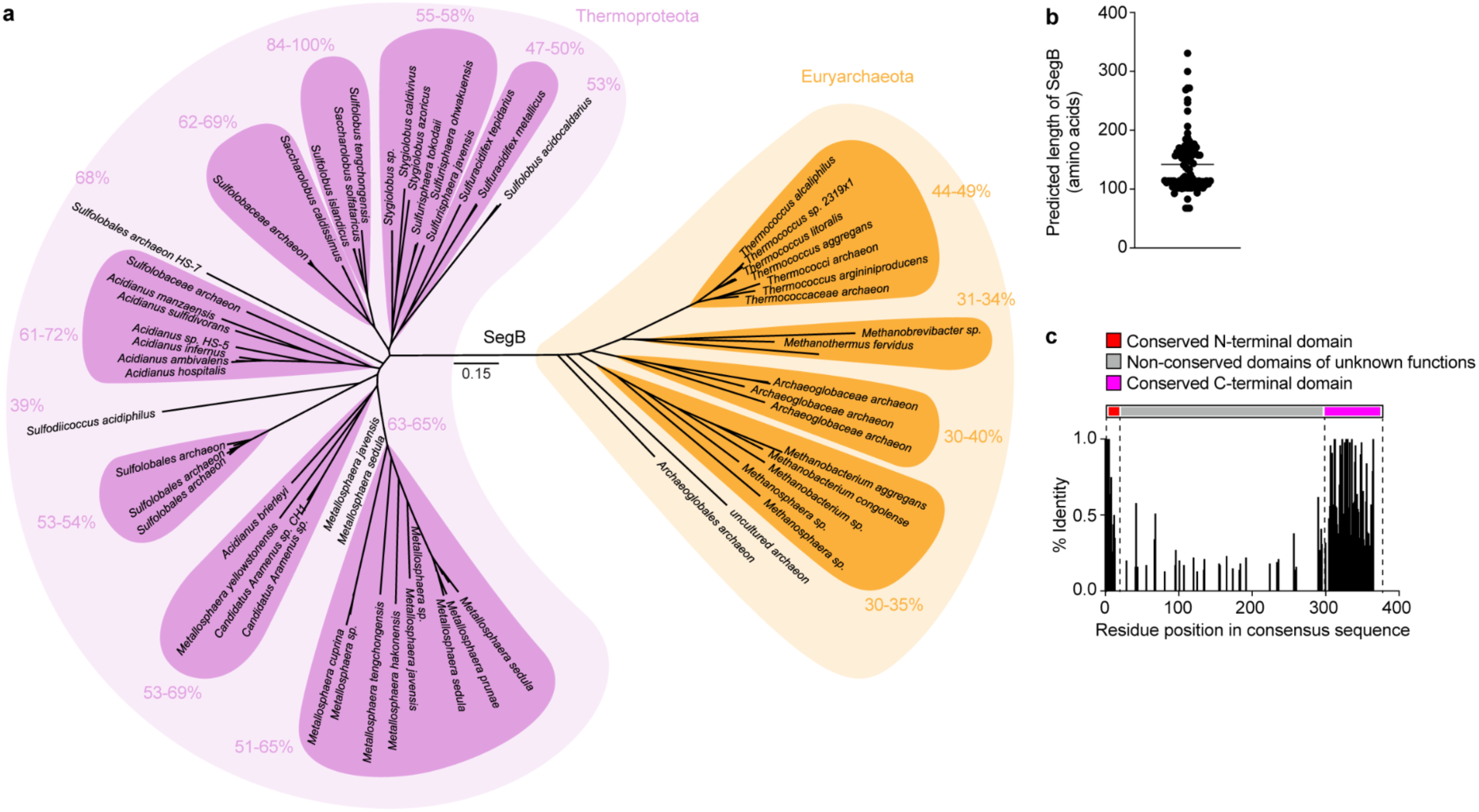
Distribution and conservation of archaeal SegB homologs. **a.** Phylogenetic tree of all archaeal SegB homologs identified. Numbers indicate sequence identity between selected sequence clusters and Sso SegB. **b.** Predicted amino acid length of all archaeal SegB homologs. **c.** Frequency of residue conservation in all archaeal SegB homologs compared to a consensus SegB sequence.

**Figure S5.**
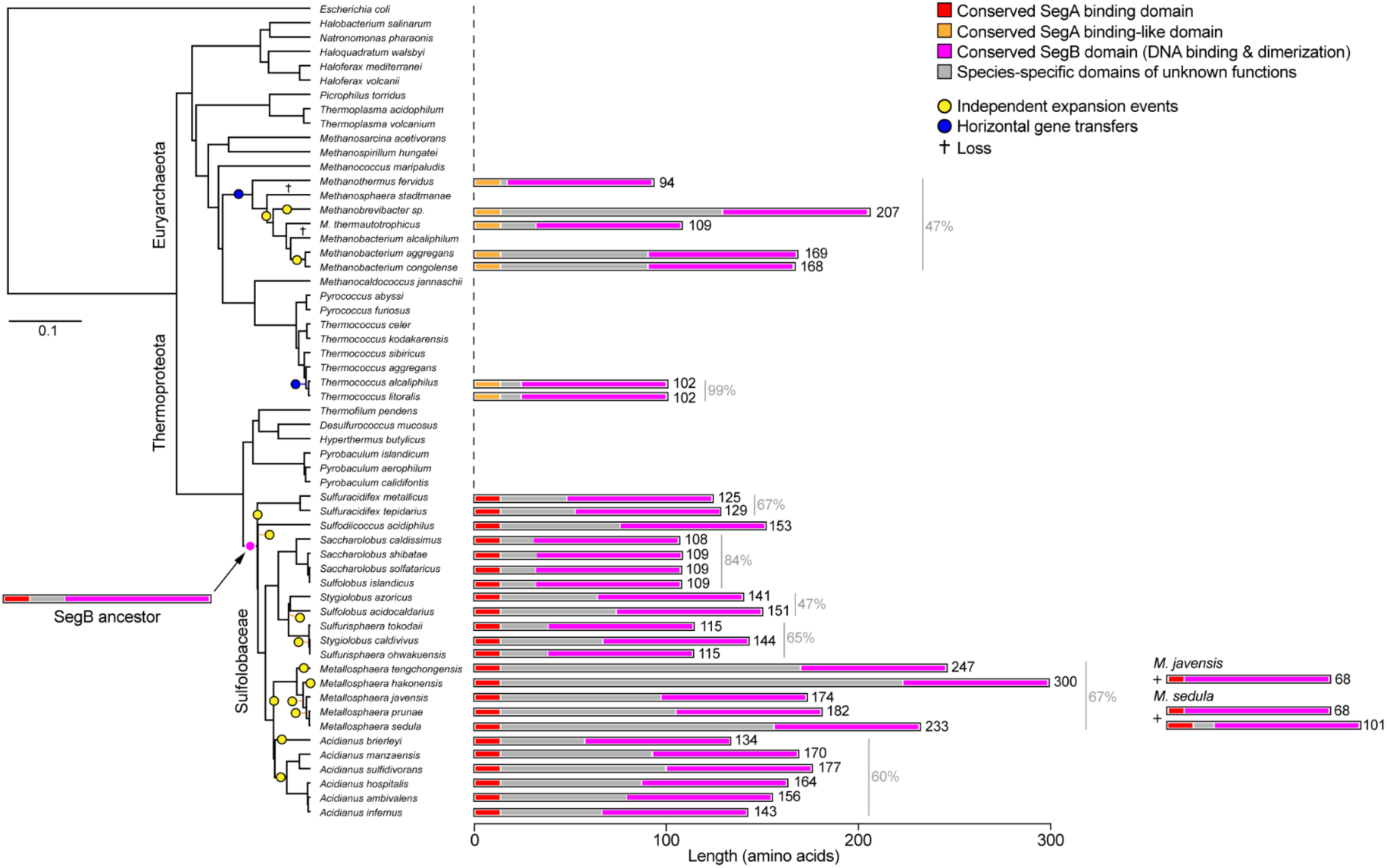
Mechanism for SegB sequence expansion and evolution of archaeal SegB. Mapping of SegB homolog lengths onto a phylogenetic tree of Thermoproteota and Euryarchaeota suggests that the ancestor of all SegB was likely a short homolog that was present in the ancestor of all Sulfolobaceae. At least eight independent occurrences of sequence expansion may have given rise to all SegB length variability in a lineage-specific manner (yellow circles). The presence of SegB in Euryarchaeota likely results from at least two horizontal gene transfers from Thermoproteota (blue circles). Numbers indicate sequence identity in selected SegB clusters.

**Figure S6.**
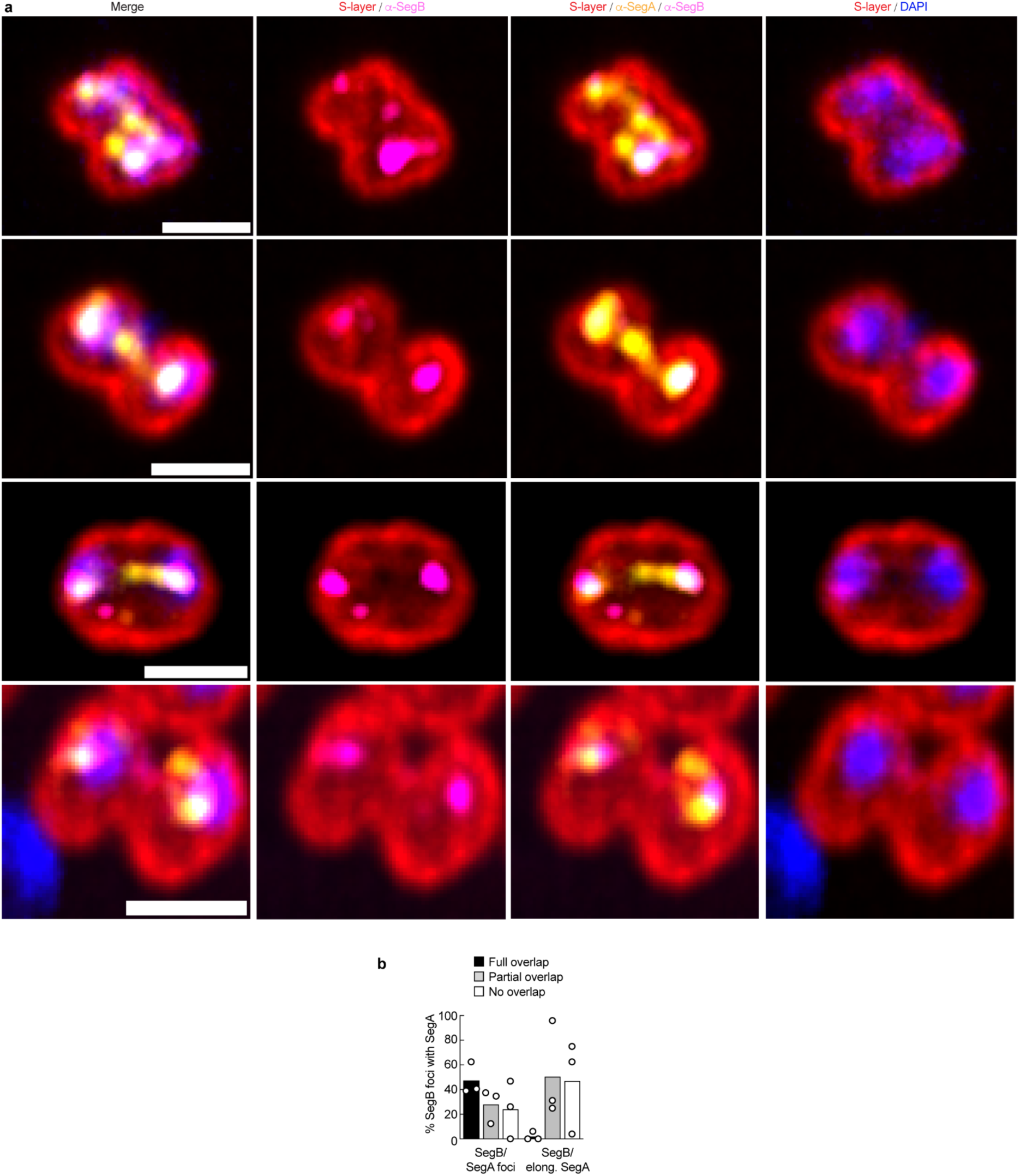
Subcellular localizations of SegA and SegB in dividing *S. acidocaldarius* (continued). **a.** Additional example immunofluorescence images of *Saci* cells with the S-layer, SegA, SegB, and DNA labeled. SegA forms multiple cellular distributions depending on the extent of division. Earlier in division (first and third row), SegA often forms puncta or extended structures near the division plane. Later in division (second and last rows), SegA tends to colocalize tightly with SegB and the compacted DNA. Scale bars are 1 μm. **b.** Quantification of the colocalization between SegB and SegA foci, or between SegB and elongated SegA figures.

## Notes

### Competing Interest Statement

The authors have declared no competing interest.

### Summary of Updates

This version of the manuscript has been revised to include X-ray diffraction data and refinement statistics.

